# Heterosis counteracts hybrid breakdown to forestall speciation by parallel natural selection

**DOI:** 10.1101/2022.01.20.477157

**Authors:** Ken A. Thompson, Dolph Schluter

**Affiliations:** Department of Zoology & Biodiversity Research Centre, University of British Columbia, Vancouver, BC, Canada; Department of Biology - Stanford University & Department of Plant Biology - Carnegie Institution for Science, Stanford, CA, USA

**Keywords:** *Gasterosteus aculeatus*, hybrid incompatibility, post-zygotic isolation, mutation-order speciation

## Abstract

In contrast to ecological speciation, where reproductive isolation evolves as a consequence of divergent natural selection, speciation by parallel natural selection has been less thoroughly studied. To test whether parallel evolution drives speciation, we leveraged the repeated evolution of benthic and limnetic ecotypes of threespine stickleback fish and estimated fitness for pure crosses and within-ecotype hybrids in semi-natural ponds and in laboratory aquaria. In ponds, we detected hybrid breakdown in both ecotypes but this was counterbalanced by heterosis and the strength of post-zygotic isolation was nil. In aquaria, we detected heterosis only in limnetic crosses and breakdown in neither ecotype, suggesting that hybrid incompatibilities are environment-dependent for both ecotypes and that heterosis is environment-dependent in benthic crosses. Heterosis and breakdown were 3× greater in limnetic crosses than in benthic crosses, contrasting the prediction that the fitness consequences of hybridization should be greater in crosses among more derived ecotypes. Consistent with a primary role for stochastic processes, patterns differed among crosses between populations from different lakes. Yet, we observed qualitatively similar patterns of heterosis and hybrid breakdown in benthic crosses and limnetic crosses when averaging the lake pairs, suggesting that the outcome of hybridization is repeatable in a general sense.

---

Ecological speciation is the process by which reproductive isolating barriers between lineages evolve as a consequence of adaptation via divergent natural selection between niches or environments (Nosil, 2012; Rundle and Nosil, 2005; Schluter, 2000, 2001). Divergent selection can lead to the evolution of post-zygotic isolating barriers when the alleles underlying adaptation to divergent environments function poorly together when combined in hybrids (Coyne and Orr, 2004). In many examples of ecological speciation, the fitness of hybrids depends critically on the ecological context (Nosil, 2012; Schluter, 2000). Hybrids formed between ecologically divergent species can have low fitness in the field because there is no niche suited to their intermediate or mismatched phenotypes (Hatfield and Schluter, 1999; Matsubayashi et al., 2010; Melo et al., 2014; Thompson et al., 2021; Zhang et al., 2021), even though they may have high fitness in benign environments (Hatfield and Schluter, 1999; Johansen-Morris and Latta, 2006). The study of ecological speciation has substantially clarified the role of ecology in driving the origin of species.

Compared to speciation by divergent natural selection, speciation by parallel natural selection—where the same phenotype is favoured by selection in different populations—has been the subject of fewer studies. Studying speciation by parallel selection is important because it can tell us about the role of adaptation by natural selection *per se*, compared to adaptation via divergent natural selection specifically, in speciation (Langerhans and Riesch, 2013). Speciation by parallel natural selection can occur via ‘mutation-order’ processes, wherein alternative alleles are fixed in populations adapting under similar selection pressures (Mani and Clarke, 1990). These alternative alleles might be incompatible when combined in hybrids and reduce their fitness (Schluter, 2009). Incompatibility might manifest in F_1_ hybrids if there is nonadditivity (Melo et al., 2019). Alternatively, incompatibility can manifest when additive alleles—which can be fixed from standing variation or *de novo* mutation—segregate in recombinant hybrids (e.g., the F_2_ generation), causing some hybrids to express transgressive phenotypes that are maladaptive in the parental niche (Barton, 2001; Chevin et al., 2014; Thompson et al., 2019; Yamaguchi and Otto, 2020). The reduction in hybrid fitness caused by the segregation of incompatible alleles in recombinant hybrids is termed ‘hybrid breakdown’ (Barton, 2001).

To test hypotheses about the evolution of post-zygotic reproductive isolation by parallel natural selection, we leveraged the unique biology of threespine stickleback (*Gasterosteus aculeatus* L.) species pairs (Fig. 1A). The species pairs are independently-derived (Jones et al., 2012; Taylor and McPhail, 1999; Wang, 2018) and occur in three lakes along the south coast of British Columbia, Canada (Fig. 1B). Each lake contains two reproductively isolated stickleback ecotypes—the benthic ecotype is large-bodied and uses high suction to feed on macroinvertebrates in the substrate and attached to vegetation, whereas the limnetic ecotype is relatively small-bodied and uses rapid jaw protrusion to feed on zooplankton in the open water (McGee et al., 2015; McGee and Wainwright, 2013; McPhail, 1984, 1992; Schluter, 1993, 1995; Schluter and McPhail, 1992). Hybrids between benthic and limnetic ecotypes are viable and fertile when raised in aquaria, but suffer fitness disadvantages under field conditions (Hatfield and Schluter, 1999; McPhail, 1984, 1992; Rundle, 2002). Among fishes, the benthic-limnetic species pairs represent one of the clearest examples of parallel phenotypic evolution (Oke et al. 2017; see also Conte et al. 2015 for detailed discussion of non-parallel aspects). The species pairs originated within the last 15,000 years after anadromous stickleback colonized newly-formed post-glacial lakes (Schluter, 1996), and are thought to have arisen through a process of ‘double invasion’, wherein the original anadromous colonists became the benthic population and a second anadromous immigrant population evolved to become the limnetic population.

**Fig. 1.**
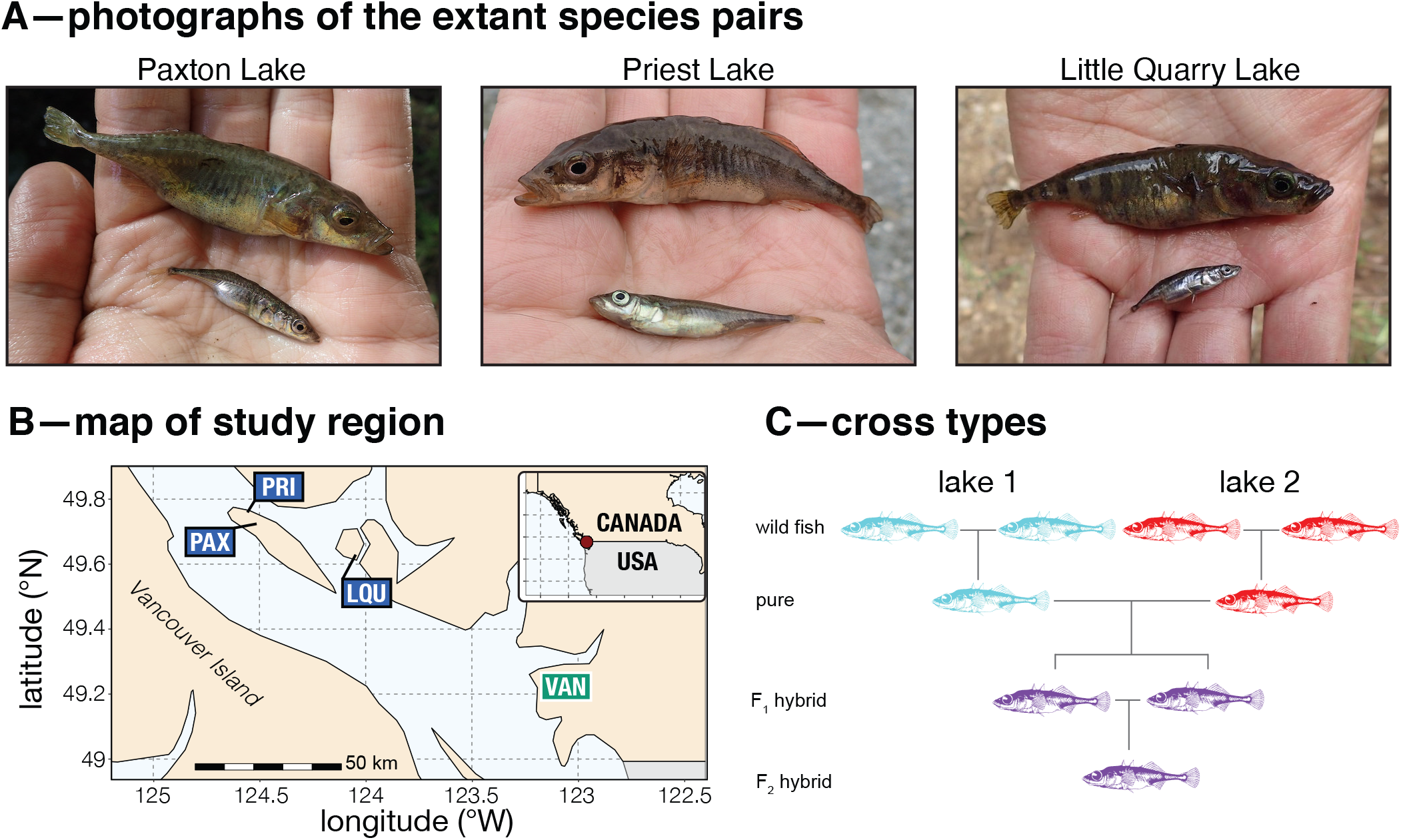
System photograph, map, and cross schematic. Panel A shows photographs of the three extant species pairs, Paxton Lake (PAX), Priest Lake (PST) and Little Quarry Lake (LQU). Each photograph was taken in 2017 and shows females of the benthic (top) and limnetic (bottom) ecotypes. Panel B shows a map of the study region and locations of the lakes (blue boxes) and the approximate location of our field and lab experiments (VAN). Panel C isa schematic illustrating the cross types made within a single ecotype for a single lake pair. (Note that no full siblings were crossed.)

We conducted experiments in the field and lab to test the fitness consequences of hybridization after parallel phenotypic evolution in the stickleback species pairs. Our experiments tracked the survival and growth of individual juvenile fish to estimate fitness. Because all known patterns of post-zygotic isolation in stickleback species pairs are extrinsic, we expected fitness differences to manifest mainly under field conditions. We expected that any of three main patterns could result. The first is that hybrids have no reduction in fitness if the genetic basis of parallel phenotypic evolution is also highly parallel, or if divergent alleles do not interact negatively. The second is hybrid breakdown, wherein opposite-ancestry divergent alleles reduce fitness when combined in hybrids. The third possibility is heterosis, wherein heterozygosity is favoured by selection. Prior research indicates a largely additive genetic basis of phenotypic evolution in the species pairs (Arnegard et al., 2014; Miller et al., 2014) that is only partially shared among different populations of the same ecotype (Conte et al., 2015). Given this, we expected that F_1_ hybrids would have similar fitness to pure crosses and that F_2_ hybrids might exhibit hybrid breakdown due to the segregation of population-specific QTL (Chevin et al., 2014; Munzur and Thompson, 2021; Thompson et al., 2019). Theory predicts that hybrid breakdown is proportional to the magnitude of evolutionary change in parallel from the common ancestor (Barton, 2001; Chevin et al., 2014). Because benthic populations are more phenotypically and genomically distant from the marine ancestral form (Chhina et al., 2022; Jones et al., 2012; Wang, 2018), we predicted that hybrid breakdown would be greater in crosses between benthic populations from different lakes than in crosses between limnetic populations from different lakes. Our results provide novel insights into the efficacy and predictability of speciation by parallel natural selection.

## Methods

### Experimental crosses

We made all possible pairwise within-ecotype crosses using the three extant benthic-limnetic species pairs—Paxton, Priest, and Little Quarry Lakes (Fig. 1C). We made pure within-population crosses (i.e., ‘parental’ or ‘non-hybrid’ crosses) for each ecotype and lake. We also generated F_1_ and F_2_ within-ecotype hybrids. All crosses were between unrelated families, but some families and individuals were used as parents more than once in separate crosses. Parents of all crosses hatched in the lab, and no wild progenitors were collected before 2016. Crosses were made in March 2020, and fish were raised in 100L aquaria until mid-June 2020 when we began the pond experiment. The data underlying the results are drawn from 111 unique crosses.

### Pond experiment

The pond experiment occurred in three semi-natural experimental ponds on the campus of the University of British Columbia. The ponds were established between 2008 and 2010 and are 25 m long × 15 m wide. The 25 m length of each pond is made up of a 12.5 m gradually sloping shallow littoral zone with macroalgae and macrophytes and a 6 m deep open-water zone (see extended data Fig. 1 from Arnegard et al. 2014 for a detailed description and diagram). Previous research indicated that the diets of benthic and limnetic fish in the ponds were similar to their diets in natural lakes (Arnegard et al., 2014). Except for their use in previous experiments, the ponds are unmanipulated environments. Each pond contained fish with ancestry from only two lakes (i.e., one lake pair; pond 4—Paxton & Little Quarry; pond 9—Priest & Little Quarry; pond 19—Priest & Paxton).

Our pond experiment included over 3,700 individual fish—at least 600 from each ecotype in each of the three ponds. Sample sizes were determined using power analyses conducted in advance of the experiment (see Supplementary Text). For crosses of both ecotypes, we introduced approximately 100 individuals of both pure crosses, 200 F_1_ hybrids, and 200 F_2_ hybrids. Total numbers of introduced fish are given in Table S1. For a visual depiction of the experimental design, see Fig. S1.

Before introduction, each fish was weighed to the nearest 0.01 g and then a sequential coded wire tag (hereafter ‘tags’; Northwest Marine Technology, Anacortes, WA, USA) was implanted into the dorsal musculature on the right side of the body. Tags can be recovered from recaptured fish at the end of an experiment to identify an individual via dissection and the use of a microscope. Detailed methodology is given the Supplementary Methods. The mean initial body mass (± SD; all cross types) was 0.56 ± 0.16 g for introduced benthic crosses and 0.44 ± 0.11 g for introduced limnetic crosses, which corresponds to approx. 25–35 mm standard length for crosses of both ecotypes (adult mass of benthic and limnetic fish is typically *>* 2 g and *>* 1 g, respectively; see Results). After tagging, fish were returned to their original tank for a 48 hr recovery period, then transported to the ponds in coolers containing their original tank water, which upon arrival at the ponds was diluted 50:50 with pond water. Fish were kept in these coolers overnight—with two aerating air stones, several plastic plants, and sections of PVC pipe—and released into the ponds the following morning. The few fish that perished before introduction had their tags extracted and read so they could be excluded from the analysis. For each pond, tagging and release occurred over approximately two weeks (release windows—Pond 4: June 14–28; Pond 9: June 29–July 10; Pond 19: July 11–24).

We retrieved surviving fish from each pond using minnow traps and by dip-netting beginning on 14 September 2020, after the experiment had run for approximately three months. Minnow traps were baited using old cheddar cheese wrapped in several layers of lightly perforated aluminum foil. After several days of trapping and netting, when fish returns slowed to fewer than five per evening of trapping, we added 2 L of 5% rotenone to each pond. The rotenone caused remaining fish to swim to the surface of the pond, where they were easily collected with a dip net. After collection, fish were euthanized with an overdose of MS-222. We recorded the fresh mass of each fish, took a photograph, and then immediately stored the fish at −20° C in 15 mL centrifuge tubes containing unique paper labels. Tags were extracted from frozen fish after lightly thawing them, then read with a microscope (Magniviewer, Northwest Marine Technology). Tags were matched with the original data to identify each fish. From this, we could determine the survival and total growth of each individual released into ponds.

### Lab experiment

While the pond experiment was ongoing, we conducted a similar experiment in the lab using siblings or relatives of fish released into ponds. The goal of this experiment was to provide a point of comparison to the pond data that could allow us to make inferences about whether the patterns observed in ponds were intrinsic or extrinsic.

The laboratory experiment was conducted using 60 × 110 L aquaria in a common recirculating system. Most tanks contained both benthic and limnetic individuals and all individuals from a given ecotype inhabiting the same tank were from the same family. 20 g of fish were added to each tank to standardize initial mass for a total of 2,151 fish. We recorded the mass of each fish at the experiment onset, but individual fish were not tagged due to logistical constraints. Immediately after we ended the pond experiment, we euthanized all surviving fish in aquaria with an overdose of MS-222, then recorded their ecotype (easily distinguished by eye) and mass (mean number of days in aquaria = 74). We estimate growth only in tanks in which all fish survived to the end of the experiment. In these tanks, we assume that—within each ecotype—the individuals’ rank order of sizes did not change between the beginning and end of the experiment and quantified the difference in mass between the initial and final sampling points as ‘growth’. We measured survival by comparing the number of fish of each ecotype in tanks between the experiment’s start and end.

### Data analysis

We fitted linear or generalized linear models depending on the fitness component. Response variables were either survival (binary; generalized linear model) or final mass (continuous; linear model). All mass variables were log-transformed (ln[*x* + 1]) prior to analysis to reduce heteroskedasticity. When comparing cross types (i.e., pure, F_1_, F_2_), we fitted models for the benthic and limnetic ecotypes separately because of significant differences in survival and body size, but use parameter estimates to calculate a composite fitness metric that we compare directly between ecotypes (see below). We analyzed pond and aquarium data separately.

Most models fitted to pond data included ‘lake pair’ as a random effect, which includes the combined effects of pond and lake pair (‘pair-as-random’ models). These models included cross type as a fixed effect with three factor levels: pure, F_1_ hybrid, and F_2_ hybrid. Initial mass was a covariate in all models. For models of growth, we included duration—the number of days between the introduction into ponds or aquaria and the final mass measurement—as a covariate. Survival models also included a duration variable: the minimum observation period, calculated as the difference between the day a fish was introduced into the pond or tank and the first day of trapping of that pond (or tank sampling day) at the end of the experiment (we do not know on which day an unrecovered fish died). These models estimate survival and growth for both ecotypes. Although we do analyze the model output directly, we base our conclusions primarily on a composite fitness metric (described below) combining the model-estimated patterns of survival and growth. These pair-as-random models thus estimate the fitness consequences of hybridization for both ecotypes across the three lake pairs.

Compared to cases of divergent evolution, the fitness consequences of hybridization after parallel phenotypic evolution are more likely to be governed by chance events during evolution, such as founder effects and the mutation-order process. It is therefore of interest to test whether the fitness consequences of hybridization differ among crosses between different lake pairs (e.g., Paxton × Priest vs. Paxton × Little Quarry). To investigate the effects of lake pair we therefore fit models to pond data that included lake pair and the interaction between cross type and lake pair as fixed effects (‘pair-as-fixed’ models). This was done even though the effect of lake pair is confounded by effects that could be specific to a single pond. In the Discussion, we address differences among the lake pairs and the possibility that some differences are present beyond pond-specific effects. These models estimate the mean survival and growth for each cross type for each lake pair for both ecotypes, which are used to estimate a composite metric of fitness, as above. This composite fitness metric is compared between the benthic and limnetic ecotypes in the main text and we present analyses of survival and growth in the supplement. In sum, these pair-as-fixed models allowed us to estimate the extent to which patterns of hybrid fitness were not repeatable among lake pairs.

We did not include family (i.e., unique fertilized clutch) or rearing tank (for large families that were split into multiple tanks) as random effects in our analysis and treat each fish as an experimental unit. Because many families were split between two or more tanks, the variance among tanks includes the variance among families. Family accounted for no additional variance when tank was included in a model estimating final mass (model comparison ANOVA; *P* = 0.24). All fish from a tank went into a pond at the same time, and the time of introduction of each tank was determined systematically to ensure maximal variation among cross types and ecotypes. We therefore account for these effects by including initial mass and duration (determined by introduction date which is the same for all fish in a tank) in all models fitted to pond data.

All analyses were done in R version 4.0.3. Mixed models were fit with the `lme4` package (Bates et al., 2014) and analyzed with ‘anova’ after loading the `lmerTest` package (Kuznetsova et al., 2014). The significance of main effects in mixed models was evaluated using the Kenward-Roger approximation for the denominator degrees of freedom (Kenward and Roger, 1997). Post-hoc analyses were done using the ‘emmeans’ and ‘pairs’ functions in `emmeans` (Lenth et al., 2020), with Tukey HSD–corrected *P*-values. Model fits were visualized with `visreg` (Breheny and Burchett, 2017). Data processing used functions in the `tidyverse` (Wickham, 2017).

### Composite fitness metric

We generated a composite fitness metric as the product of survival and growth (DiVittorio et al., 2021; Patton et al., 2021). Because survival and growth cumulatively affect fitness—a group with ½ the survival and ½ the fecundity of another would have a relative fitness of ¼—composite fitness metrics more accurately capture group differences than do either survival or growth alone. The conclusions of analyses with multiplicative fitness are more conservative (i.e., smaller differences among groups) than an alternative analysis that estimated fitness using more assumption-laden estimates of fecundity and overwinter survival (Table S2). We estimated fitness of each hybrid cross type and combined pure parents in each pond as the product of estimates of survival and growth from our fitted models. Mass estimates were back-transformed from the log scale for fitness calculations (Lande and Arnold, 1983). We divided each cross type’s fitness value by the composite fitness of the pure crosses to standardize them. Thus, F_1_ and F_2_ fitness is relative to the pure crosses, which have a composite fitness value of 1.

We used bootstrapping to compare composite fitness among cross types. In each bootstrap iteration, we resampled survival and growth of individuals of each cross type (i.e., pure [parents], F_1_ hybrid, F_2_ hybrids) so that the sample sizes were identical to the authentic dataset. For the pond experiment, resampling was conducted within the individual aquaria that were tagged and released into the ponds because variation in tagging and release time was controlled systematically during the experiment—that is, resampling within ‘tank’ is necessary to keep the value of the ‘duration’ covariate consistent across bootstrap iterations. For consistency, resampling for the aquarium experiment also conducted within individual aquaria. We ran identical models as conducted above, with lake pair as a random effect where appropriate. Because we only had three lake pairs, sample sizes were too low to resample levels of the ‘pair’ random effect. Each iteration returned a point estimate for fitness as the product of the estimated survival and growth values for each cross type for both ecotypes. To compute confidence intervals of composite fitness, the bootstrapped fitness estimates were divided by the composite fitness of pure crosses observed in the authentic data (for that same ecotype). Alternative bootstrap resampling procedures were similar or less conservative (Fig. S2).

We considered a comparison of composite fitness to be statistically significant if the 95% bootstrap confidence interval of the difference being estimated did not include 0. We tested all pairwise differences among pure, F_1_ hybrid, and F_2_ hybrid crosses. The pure – F_1_ difference tests for heterosis. We tested for hybrid breakdown by determining if the fitness of F_2_ hybrids was less than the mid-point of pure crosses & F_1_ hybrids. We compute post-zygotic isolation as the difference between pure crosses and F_2_ hybrids, and only interpret data from ponds for this metric.

We also used bootstrapping to compare the magnitude of heterosis and breakdown between benthic and limnetic ecotypes, and between pond and aquarium studies within both ecotypes. We first extracted the magnitude of the composite fitness differences between relevant groups (heterosis = F_1_–pure; hybrid breakdown = F_2_ –[pure:F_1_ midpoint]) which we term the ‘magnitude of [heterosis or hybrid breakdown]’. We computed the difference in these resampled magnitudes between benthic and limnetic crosses within each bootstrap iteration and then tested whether the bootstrap confidence intervals included zero. To compare aquarium and pond data, a similar procedure was used except the difference was computed for each iteration of the pond- and aquarium-based estimates of heterosis and breakdown.

## Results

### Summary of pond experiment

We recovered and read tags from 59.6% of fish that were introduced into ponds (number released = 3,713, number recaptured = 2,213). Across the entire dataset, the initial mass of a fish and the number of days it spent in the ponds were both positively correlated with final mass (Fig. S3 & S4). Recapture rate, which we assume reflects survival, was 78.2% for benthic crosses and 40.9% for limnetic crosses (‘ecotype’ main effect: *χ*^2^(1) = 402.92; *P* < 0.0001). Benthic fish accumulated 0.98 ± 0.018 g [SE] greater body mass than limnetic fish (*F*_1,2191.1_ = 2808.4; *P* < 0.0001). Recaptured benthic fish were on average 3.8× their initial mass and recaptured limnetic fish were on average 2.4× their initial mass.

### Relationships among cross types

For both benthic and limnetic ecotypes, cross type (i.e., pure, F_1_, and F_2_) often had a significant effect on survival and growth in ponds (Fig. 2; pair-as-random models). Growth—but not survival differed among benthic cross types (main effects; survival—*χ*^2^(2) = 1.β1, *P* = 0.45, growth—*F*_2,437_ = 8.72, *P* = 0.0002), and both survival and growth differed among limnetic cross types (main effects; survival—*χ*^2^(2) = 7.86, *P* = 0.02; growth— *F*_2,744_ = 17.56, *P* < 0.0001). In all cases where significant differences were detected among cross types in ponds, the pattern was F_1_ hybrid *>* pure = F_2_ hybrid, with the F_2_ hybrid mean being slightly lower than the mean of pure crosses (though statistically equivalent). As in the ponds, in aquaria we detected growth but not survival differences among benthic cross types (Fig. S5A; survival—*χ*^2^(2) = 2.88; *P* < 0.23; growth—*F*_2,432_ = 50.75; *P* < 0.0001) and detected differences among limnetic cross types in both survival and growth (Fig. S5B; survival—*χ*^2^(2) = 19.02; *P* < 0.0001; growth—*F*_2,258_ = 50.75; *P* < 0.0001). Where differences were detected in aquaria, patterns for benthic crosses were pure *>* F_1_ hybrid = F_2_ hybrid while for limnetic crosses the pattern was pure < F_1_ hybrid = F_2_ hybrid.

**Fig. 2.**
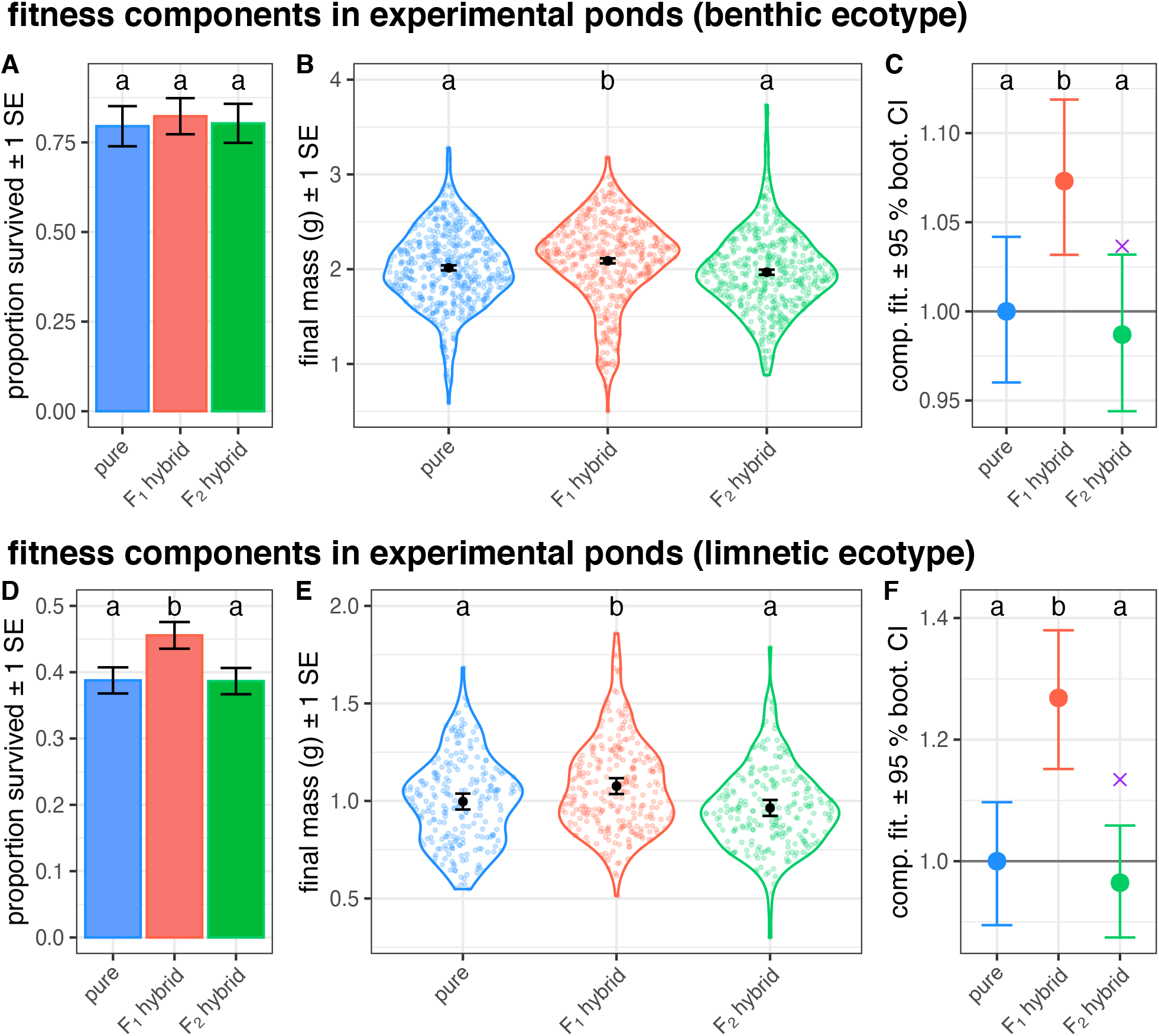
Survival, growth, and composite fitness of pure population crosses and their F_1_ & F_2_ hybrids in the experimental ponds (pair-as-random models). Data for benthic crosses are presented in panels A–C and data for limnetic crosses are shown in panels D–F. Survival and growth plots show estimated marginal means (Lenth et al., 2020) and standard errors. Mean survival (A & D) was estimated using generalized linear mixed models. Growth panels (B & E) show partial residuals of individual-level variation in violin plots for final mass—points are individual fish. Composite fitness (C & F) was estimated by multiplying survival and growth and dividing by the pure value, and 95% bootstrap confidence intervals generated by resampling the data and re-running the models. Different letters indicate a difference at *P* = 0.05 from models (survival and growth) or bootstrap confidence intervals (fitness). Purple ‘×’ symbols indicate the pure:F_1_ hybrid midpoint as a reference for the expected F_2_ mean in the absence of hybrid breakdown.

The remainder of our analyses use the composite fitness metric because the differences among groups are similar for our composite fitness metric as they are for survival and growth (survival and growth patterns are qualitatively similar to one another for both ecotypes). All *P*-values are from bootstrap resampling.

We detected significant heterosis—defined as a difference in fitness between pure crosses and F_1_ hybrids—in both ecotypes in ponds (Fig. S6; pair-as-random models). Benthic F_1_ hybrids had 7% greater fitness than pure benthic crosses (*P* = 0.012), and limnetic F_1_ hybrids had 27% greater fitness than pure limnetic crosses (*P* = 0.002). This 3.9× greater heterosis in limnetic F_1_ hybrids compared to benthic F_1_ hybrids represents a significant difference in the magnitude of heterosis between ecotypes (Fig. S7; *P* = 0.032). In aquaria, we observed significant heterosis in limnetic crosses, with F_1_ hybrids having 19% greater fitness than pure crosses (*P* = < 0.001) whereas composite fitness was reduced by 16% in benthic F_1_ hybrids compared to pure crosses (*P* < 0.001). The magnitude of heterosis was significantly greater in ponds than in aquaria for benthic F_1_ hybrids (*P* = 0.024), but did not differ between experiments for limnetic F_1_ hybrids (Fig. S8; *P* = 0.68).

We also detected significant hybrid breakdown— defined as a difference in fitness between F_2_ hybrids and the midpoint of pure crosses and F_1_ hybrids—in both ecotypes in ponds (Fig. S6; pair-as-random models). The composite fitness of benthic F_2_ hybrids was reduced by 5% below the pure:F_1_ hybrid midpoint (*P* = 0.042), and the composite fitness of limnetic F_2_ hybrids was reduced by 16% below pure:F_1_ hybrid midpoint (*P* = 0.016). Similar to heterosis, the magnitude of breakdown was (3.1×) greater among limnetic crosses than among benthic crosses, however this difference did not meet the threshold for statistical significance (Fig. S7; *P* = 0.08). We did not detect hybrid breakdown in either ecotype within aquaria because observed F_2_ fitness was slightly greater than F_1_ fitness and the two were statistically indistinguishable. As a result, the magnitude of hybrid breakdown was significantly greater in ponds than in aquaria for both ecotypes (Fig. S8; benthic—*P* = 0.024; limnetic—*P* < 0.001).

We detected no evidence of post-zygotic isolation in ponds in the pair-as-random models. Specifically, for both the benthic and limnetic ecotype, the composite fitness of pure crosses and F_2_ hybrids was statistically indistinguishable (benthic—*P* = 0.68; limnetic—*P* = 0.63).

### Variation among lake pairs in ponds

Patterns differed among the three lake pair crosses (Fig. 4; see Fig. S9 for underlying fitness components). In benthic crosses, we detected heterosis in one of the three lake pairs (Paxton × Priest [*P* < 0.001]) and detected breakdown in two of the three lake pair crosses (Paxton × Priest [*P* = 0.002] & Little Quarry × Priest [*P* = 0.014]). In limnetic crosses, we detected heterosis in two of three lake pairs (Paxton × Priest [*P* = 0.01] & Little Quarry × Paxton [*P* < 0.001]) and breakdown in one of the three lake pairs (Little Quarry × Paxton [*P* < 0.001]; breakdown approached significance in Paxton × Priest cross [*P* = 0.054]).

**Fig. 3.**
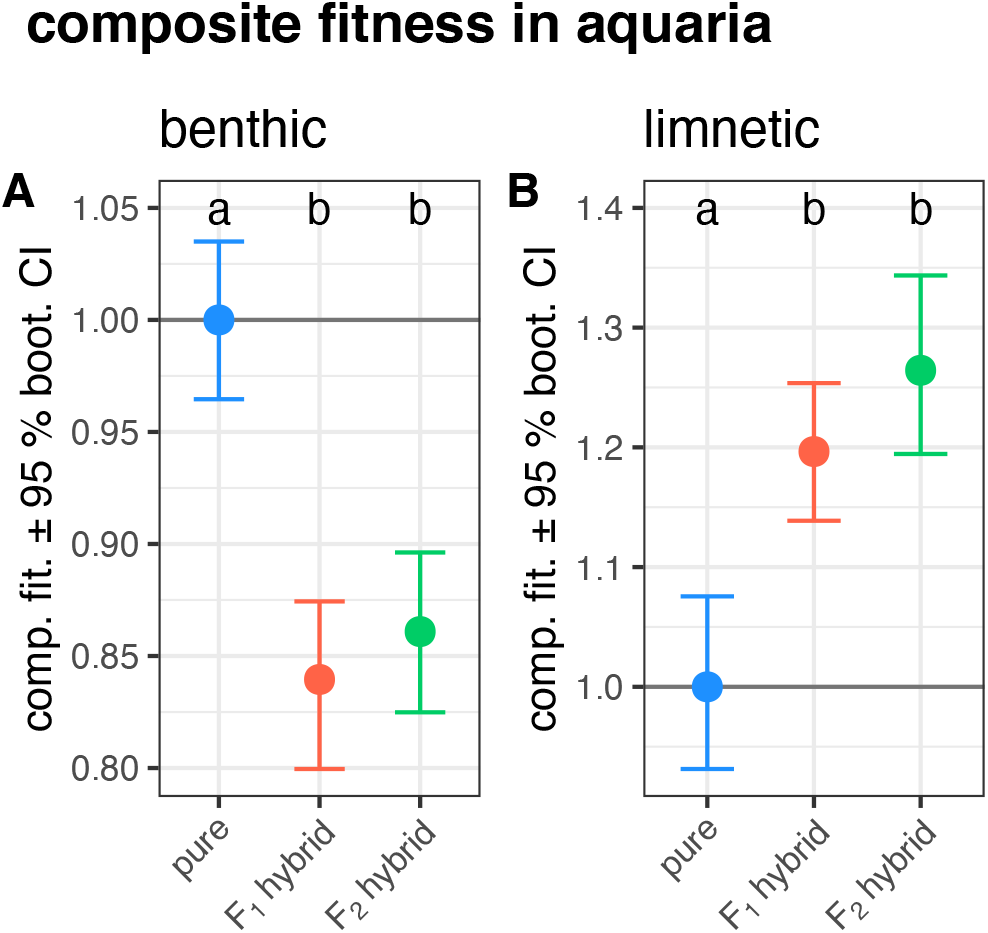
Composite fitness among (A) benthic and (B) limnetic cross types in aquaria. Panels show the observed composite fitness (points) and 95% bootstrap confidence limits. The underlying estimates of survival and growth are shown in Fig. S5. Letters indicate differences at *P* = 0.05.

**Fig. 4.**
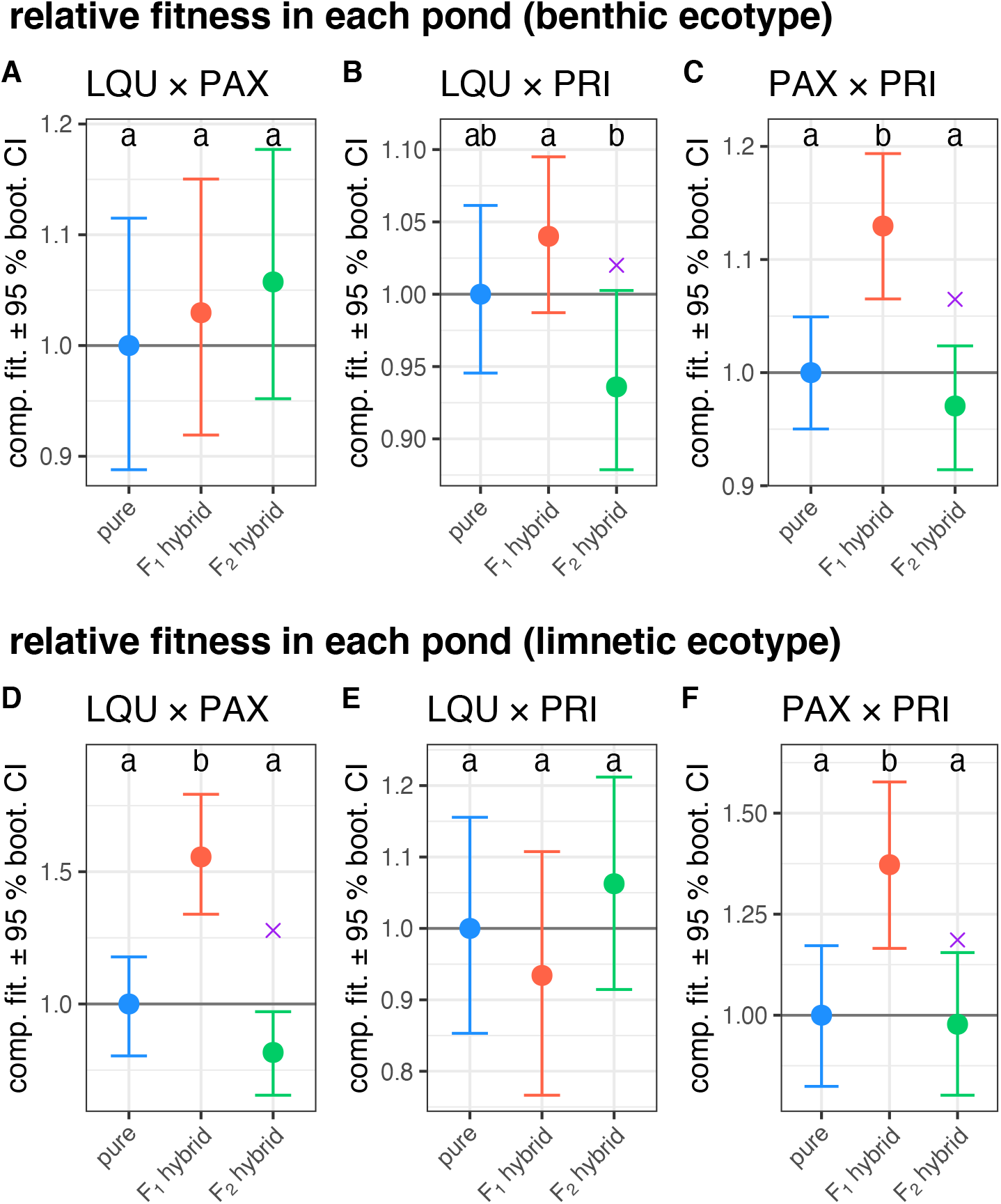
The fitness consequences of hybridization differed among lake pairs (pair-as-fixed models). The upper panels show patterns in crosses between three pairs of benthic populations and the lower panels show results of crosses between three limnetic pairs. Points are estimates from our statistical models, and error bars are 95% bootstrap confidence intervals from 1,000 bootstrap replicates. Letters indicate differences at *P* = 0.05. Purple ‘×’ symbols indicate the pure:F_1_ hybrid midpoint as a reference for hybrid breakdown (only shown for crosses where F_2_ < F_1_).

## Discussion

In this study, we experimentally quantified the fitness consequences of hybridization after parallel phenotypic evolution in populations of the same stickleback ecotypes that have evolved repeatedly from a common ancestral form. In agreement with predictions from theoretical models of mutation-order speciation, we detected significant hybrid breakdown between benthic populations and between limnetic populations in experimental ponds. However, we also detected an equal magnitude of heterosis in ponds and as a result, the composite fitness estimates for pure crosses and F_2_ hybrids were invariably indistinguishable. We also found that the magnitude of heterosis and hybrid breakdown were more than 3× greater in limnetic crosses than in benthic crosses, which contrasts the prediction that the ecotype most divergent from the ancestor should show the greatest amount of breakdown. Data from aquarium-raised fish reveal that heterosis might be intrinsic—occurring via mechanisms that are environment-independent—in limnetic crosses but that hybrid breakdown only occurred in ponds. Finally, although patterns differed among the three lake pairs for crosses of both ecotypes (estimated via pair-as-fixed models), the overall qualitative patterns of heterosis and hybrid breakdown were similar for benthic and limnetic crosses (estimated via pair-as-random models). Below, we discuss the mechanisms that might underlie our findings and their relevance for our broad understanding of speciation by parallel natural selection.

We detected hybrid breakdown that reduced F_2_ hybrid fitness below the pure:F_1_ midpoint by 5% in benthic crosses and by 15% in limnetic crosses, which suggests that incompatible alleles segregated in the F_2_ generation (Barton, 2001). Hybrid breakdown was only detected in ponds, and thus our result contributes to a growing body of research, in stickleback and other species, suggesting that Bateson-Dobzhansky-Muller hybrid incompatibilities (Bateson 1909; Dobzhansky 1937; Muller 1942) can have environment-dependent effects on fitness (Johansen-Morris and Latta, 2006). In F_2_ benthic × limnetic hybrids, Arnegard et al. (2014) and Keagy et al. (2016) found evidence that mismatched trait combinations reduced individual fitness. By re-analyzing the genetic data from Arnegard et al. (2014) and others, we previously found evidence that a genetic signature of selection against hybrid incompatibilities was detectable in pond-raised stickleback hybrids, but not those raised in aquaria (Thompson et al., 2022). Studies in systems as diverse as yeast (Ono et al., 2017), anole lizards (Bock et al., 2021), and sunflowers (Sambatti et al., 2008) have drawn similar conclusions. In line with emerging findings that recognize ecology’s role in shaping epistasis within natural populations (Nosil et al., 2020), it is increasingly clear that ecology can also mediate the fitness consequences of hybridization between different natural populations.

In spite of significant F_2_ hybrid breakdown, we detected no post-zygotic isolation because of strong heterosis. We detected heterosis in aquarium-raised limnetic F_1_ hybrids but not in benthic F_1_ hybrids, which is consistent with previous findings that features of the environment can underlie heterosis (Wagner et al., 2021). F_1_ heterosis (and F_2_ breakdown) is regularly observed in plants (Hahn and Rieseberg, 2017; Rhode and Cruzan, 2005; Rosas et al., 2010; Vasseur et al., 2019), but is not as common in outbred wild animal species. Some models suggest that heterosis is a general property of hybrid systems (Fiévet et al., 2018), and it could be the case that ecological divergence overrides heterosis in most animal systems that have been subject to experimentation (e.g., in divergent crosses between [rather than within] benthic and limnetic ecotypes [Hatfield and Schluter 1999]). Studies of model systems often find evidence for heterosis during the early stages of divergence, followed by declines over longer timespans (Dagilis et al., 2019; Wei and Zhang, 2018). Thus, it is likely that heterosis is an ephemeral phenomenon in our system while hybrid breakdown is enduring.

Our findings are opposite to the prediction from theory that crosses among derived populations should have more exaggerated fitness consequences than crosses among ancestor-like populations. Specifically, we found that the fitness consequences of hybridization were greater in crosses between limnetic populations than in crosses between benthic populations. Greater heterosis and breakdown among limnetic crosses might best be explained by differences in the selective regimes experienced by the two ecotypes in the experimental ponds. Limnetic crosses survived at approximately half the rate of benthic crosses and so experienced greater opportunity for selection (variance in composite fitness was greater among limnetic individuals than among benthic individuals). Food was unlikely to be limiting in ponds and so we do not suspect starvation: all three experimental ponds lacked fish for at least two years prior to the experiments and pond-raised fish grew considerably larger than fish in aquaria which were fed to satiation twice daily (Fig. S11). We suspect that mortality was largely caused by enemies such as piscivorous birds—we regularly observed Belted Kingfishers (*Megaceryle alcyon*), Common Mergansers (*Mergus merganser*), and Great Blue Herons (*Ardea herodias*) near or in the ponds. Adult backswimmers (Notonectidae) and damselfly and dragonfly (Odonata) nymphs were abundant though they rarely kill stickleback in the size range (25–35 mm standard length) that we introduced (Marchinko, 2009; Reimchen, 1980). Several fish were found dead during the experiment with large (approx. 5 mm) wounds on their abdomens, and several fish were recaptured at the end of the experiment with similar wounds that had healed. In addition, several fish had worm-like parasites that were visible through their skin. Differences in growth among cross types, which generally showed similar qualitative patterns to differences in survival, might have resulted from particular genotypes being less able to capture evasive prey. Greater opportunity for selection in limnetic crosses could therefore have resulted in the larger magnitudes of heterosis and hybrid breakdown compared to benthic crosses. More broadly, our findings imply that the fitness consequences of hybridization might de-part from theoretical expectations based on the ecology of the group being studied.

Although patterns of heterosis and hybrid breakdown varied among the three lake pairs for both benthic and limnetic crosses (Fig. 4), qualitative patterns were similar for the two ecotypes in the pair-as-random models (Fig. 2). This finding—variability and unpredictability at the level of individual lake pairs, but predictability when considering different lake pairs of the same ecotype together—is entirely consistent with theory on the process of speciation by parallel natural selection. In cases where reproductive isolation evolves as a consequence of divergent evolution, populations are expected to fix different alleles during adaptation and hybrid fitness may be largely determined by selection against intermediate or mismatched hybrid phenotypes (Hatfield and Schluter, 1999; McBride and Singer, 2010). Although the same alleles might be favoured in different populations under parallel selection, stochasticity in the origin and fixation of *de novo* mutations or differences in the founding population’s standing variation lead to differences in the extent of genetic parallelism (Fang et al., 2020; Mani and Clarke, 1990; Schluter, 2009; Thompson et al., 2019). The fitness of hybrids between phenotypically similar parent populations is largely determined by the degree of genetic parallelism during adaptation (Thompson et al., 2019). Despite this variation among lake pairs, we found evidence for counteracting heterosis and hybrid breakdown among benthic and limnetic cross types in the pair-as-random models. This finding implies that broad patterns in the genetics of parallel phenotypic evolution are shared between benthic and limnetic populations. Thus, while we should expect stochasticity at the level of individual lake pairs, predictable patterns might emerge if enough pairs are considered.

Our experiments make a number of assumptions that should be acknowledged. First, an implicit assumption is that the experimental ponds are reasonable stand-ins for natural lakes. This assumption is common to all studies taking place outside of a species’ source habitat. Previous studies have found that the diet of benthic and limnetic stickleback in these same ponds are similar to fish from their native lakes (Arnegard et al., 2014), which could alleviate this concern. Our crosses also only capture natural selection, and it is possible that sexual selection could act differently (Keagy et al., 2016). We also note that the different populations of the same ecotype are phenotypically similar but not identical (Conte et al., 2015), and also had differences in mean fitness (Fig. S10). We emphasise that we cannot decouple ‘lake pair’ effects from ‘pond’ effects because all experiment data from each lake pair come from a single pond. We suspect that the *magnitude* of differences among cross types could differ among ponds if the same lake pair was considered in multiple ponds (e.g., |F_1_ fitness – F_2_ fitness|). However, our conclusions rely mostly on the rank-order of cross types. We believe it is unlikely that the rank-order of cross types is affected by pond because this would require there to be, for example, a niche specific to F_1_ limnetic hybrids that is inaccessible to pure crosses and F_2_ hybrids. Such a scenario is difficult to conceive of. Finally, our experiment did not consider ecological selection at the larval stage (i.e., < 1 mo.), which can be extremely strong in fish (China and Holzman, 2014), nor did we consider overwinter survival (Carlson et al., 2010).

In conclusion, our study did not detect post-zygotic barriers to gene flow between independently derived populations of either the benthic or limnetic ecotype in the threespine stickleback species pairs. Pre-mating isolation is weak between allopatric populations of the same ecotype (Rundle et al., 2000) at least in part because mate choice is associated with body size (Conte and Schluter, 2013; McKinnon and Rundle, 2002; Nagel and Schluter, 1998) and shape (Bay et al., 2017)—preferences which have been strengthened by reinforcement (Rundle and Schluter, 1998). Because F_2_ hybrids had equivalent fitness to pure crosses, allopatric populations of the same ecotype would almost certainly collapse if brought together in sympatry. The observed heterosis might render a collapse even quicker (Irwin, 2020). Parallel evolution over longer timescales, where populations fix alternative alleles with substantial ‘intrinsic’ fitness effects, might be the primary pathway by which post-zygotic barriers arise via parallel natural selection (Melo et al., 2019). We conclude that there has been little progress toward speciation via parallel natural selection in this system.

## Acknowledgements

Feedback from S. Aguillon, Y. Brandvain, D. Irwin, G. Henriques, S. Otto, K. Peichel, L. Rieseberg, M. Schumer, R. Stelkens, and the Schumer and Schluter lab groups at Stanford and UBC improved the manuscript. We are grateful to J. Dafoe for providing passage to Nelson Island in 2017 & 2018, and to Jim & Arron for accommodation in 2018. The Zinn Family graciously provided us with passage to, and accommodation on, Nelson Island in 2019. G. Vander Haegen provided invaluable guidance for using CWTs. L. Alford, M. Ankenmann, J. Bizon, S. Blain, M. Boehm, L. Chavarie, A. Chhina, K. Chu, J. Dafoe, N. Frasson, C. Gerlinsky, A. Jevtic, M. Kinney, M. Mikkelsen, K. Nikiforuk, A. Munzur, M. Osmond, M. Roesti, J. Rolland, M. Urquhart-Cronish, J. Viliunas, and R. Yorque provided assistance in the lab and field. B. Gillespie, P. Tamkee, and E. Lotto provided critical support via the UBC inSEAS facility. KAT was supported by an NSERC Canada Graduate Scholarship, a UBC Four-Year Fellowship, and a British Columbia Graduate Fellowship. D.S. was supported by an NSERC Discovery Grant, a Canada Research Chair, and Genome British Columbia.

## Author contributions

Both authors planned research. KAT collected wild fish with assistance from DS, raised animals, conducted the experiments, collected & analyzed data with input from DS, and wrote the first draft of the manuscript. Both authors revised the manuscript.

## Data accessibility

All data and analysis code used in this article will be deposited in a repository (e.g., Dryad) at the time of publication. They have been made available to reviewers along with the version of this manuscript that is currently being considered for publication. If readers are interested in the underlying data or analysis code during the review period, please contact the corresponding author (KAT) who will be glad to send them.

## Supplementary methods

### Determining experiment size

Before the experiment, we conducted a power analysis to determine a sample size that would be necessary to detect a 2% difference in the mean of a fitness component between any two cross types within a pond at the 5% significance level with 80% probability. To estimate the level of variability in fitness we might observe in a pond experiment, we used the standard length data of Arnegard et al. (2014) as our fitness metric. We determined that a comparison of approximately 100 fish in the two cross types was sufficient for this purpose. We anticipated that 50% of fish would perish during the experiment—so for each pond we introduced 200 fish per cross type (i.e., pure, F_1_ hybrid, and F_2_ hybrid) per ecotype.

### Experimental animals

Wild fish were collected from Priest and Paxton Lakes (Texada Island, BC, Canada) and Little Quarry Lake (Nelson Island, BC) from 2017–2019. Two Paxton benthic males were collected from a pond population on UBC campus founded with wild Paxton Lake benthic fish in 2016. All fish in the experiment were therefore ≥ 2 generations removed from the wild. All benthics were captured using minnow traps as were most limnetic males. Gravid limnetic females were caught almost invariably by dip-netting. One Paxton limnetic family and one Little Quarry limnetic family was raised from a nest-collected clutch. In these cases we closely examined the resulting fish to ensure none were benthic × limnetic hybrids.

Wild fish were crossed either at the lakeside or at the lab by gently stripping the eggs from a gravid female fish into a small Petri dish filled with lake or aquarium water. Male parents were euthanized with an overdose of MS-222, and their testes were removed and placed in the Petri dish. A fine paintbrush was then used to release sperm from the testes and ensure it was well mixed among the eggs in the clutch. Crosses for the present experiment were all made in the lab in much the same way. Males were occasionally used to fertilize multiple clutches. Due to logistical constraints, we made crosses as females became gravid.

Crosses were made from 4 March to 9 April, 2020, and raised in 5 ppt saltwater (Instant Ocean) within 110 L aquaria with a small amount of methylene blue added as a fungicide. These two additions reduce the loss of clutches to fungus and also reduce labour required to raise healthy fish (because *Artemia* nauplii can live for 24 hr in 5 ppt saltwater). Larval fish were fed *Artemia* nauplii daily. Due to logistical constraints we could not monitor hatching success but note that we observed very little mortality. In general, there is very limited evidence that F_1_ or F_2_ hybrid stickleback exhibit ‘intrinsic’ deficiencies (Lackey and Boughman, 2017).

After the fish were large enough to handle without risk of mortality (approx. 3–4 weeks), we split large families into multiple tanks and culled excess fish. At this time we began feeding fish chopped frozen bloodworms and conducting weekly 50% water changes with fresh dechlorinated water. After splitting and culling we kept the density of fish to fewer than 30 individuals per aquarium. When fish were large enough, we added unchopped bloodworms, spirulina adult brine shrimp, and chopped mysis shrimp to their diets. Eventually, the mysis shrimp were fed whole. Fish were collected under Species at Risk Act (SARA) permits from Fisheries and Oceans Canada (SARA 16-PPAC-00004, 17-PPAC-00002, 18-PPAC-00006, 19-PPAC-00006) and the British Columbia Ministry of Forests, Lands and Natural Resource Operations (SU17-258923, MRSU18-288855, MRSU18-454239). All animal care protocols were approved by the University of British Columbia Animal Care Committee (A16-0044, A20-0050).

### Coded wire tagging

Here we fully document our methodology for using sequential coded wire tags (hereafter ‘tags’). We ordered sufficient quantities of tags from Northwest Marine Technology (https://www.nmt.us/; Anacortes, WA, USA). The tags come on sheets with two columns—one ‘fish’ column and one ‘reference’ column. Although tags appear in sequential order, the numbers do not increase in perfect 1:1 association with tag position. Because of this, the unique sequences on tags cannot be inferred without some possibility of error. It is therefore neccessary to pre-read some tags. We never used adjacent (i.e., on the same row) tags for fish bound for the same pond. This allowed us to pre-read fewer tags without risking error. Specifically, we pre-read every third tag in the ‘reference’ column using a Magniviewer (Northwest Marine Technology) and could reliably infer other tags because we know which pond each fish was retrieved from.

Before tagging, fish were anaesthetised with MS-222. We then used a single-shot CWT injector (Northwest Marine Technology) to inject tags beneath the skin on the fish’s dorsal musculature. Fish were injected while laying on their side atop a large sponge, and their head was covered with a paper towel soaked in water from their original tank. Light pressure was applied to the head and caudal peduncle to stabilize the fish during injection. Injection was easiest if the lateral plates were used as a leverage point to implant the head of the needle beneath the skin. (Limnetic fish, with their denser musculature and invariable presence of lateral plates, were easier to inject). We found maximal success when the push rod was not pushed to be maximally extended—in fact this can cause the CWT to emerge from the fish, rather an extension of about ¾ worked best. When injecting, we took care that the pectoral fin was either oriented toward the head or down toward the ventral area, so as to not be pierced by the tagging needle. Fish were kept temporarily in aerated water from their original tank for recovery and then moved back to their original tank before introduction into the ponds three days later. Methylene blue was added to their original tank to prevent infection of the tagging wound. We estimate that each fish took approximately 30 s to anaesthetize, weigh, and tag, and it took about two min to pre-read a page of tags.

At the end of the experiment we retrieved and read tags from each fish. We located tags under a dissecting microscope. Tags were always handled delicately because metal forceps can easily damage them. Most often, a scalpel was used to cut away skin and visually identify the tag. After cutting away muscle, the magnetic tag is attracted to the scalpel, where it can be gently placed onto the end of the magnetic ‘pencil’ that is used with the MagniViewer. Removed tags were read with the MagniViewer and then stored for later reference if necessary. A magnetic T-wand (NMT) was occasionally used to confirm the presence of a tag in a fish (or a piece of its flesh).

Ambiguities and possible transcription errors were identified by ensuring a match between ecotype (benthic or limnetic; assigned visually from re-captured fish) and pond, checking for duplicates and implausible values, and ensuring each recovered tag was assigned to a fish.

### Estimating fitness via fecundity and overwinter survival

In the main text we use a conservative fitness estimate that incorporates only survival and growth. A more complicated and possibly more accurate estimate of fitness would account for overwinter survival and fecundity. This estimate of composite fitness considers three components. The first, survival during the experiment (i.e., ‘summer survival), was directly measured. The next two fitness components consider the fitness effects of body size (directly measured) on overwinter survival and fecundity (both estimated). Absolute fitness was calculated as summer survival × (estimated) winter survival × (estimated) fecundity. Composite fitness is reported as the absolute fitness of each cross type divided by the cross type with the highest absolute fitness (within ecotype), and thus is relative to the pure crosses.

We estimated relative overwinter survival and fecundity using previously published data. These previous studies used standard length to predict fitness components, and we measured approximately 50 individuals of both ecotypes to estimate the mass–length relationship for both. Quadratic models had a high explanatory pattern for crosses of both ecotypes (*r*^2^_limnetic_ = 0.88; *r*^2^_benthic_ = 0.79) (Fig. S12).

To estimate relative overwinter survival, we conservatively interpret the analysis of Carlson et al. (2010), who found that standard length was positively associated with overwinter survival in Alaskan stickleback. The specific relationship between length and survival was either quadratic (concave) or linear, depending on the year (Carlson et al., 2010). We conservatively assume that relative overwinter survival is a linear function of standard length. I estimated fecundity using two previously published datasets from stickleback experiments in the UBC Experimental Ponds (Fig. S13). Schluter et al. (2021) estimated fecundity in F_2_ marine × freshwater stickleback hybrids. Specifically, the fitness of over 200 F_2_ hybrid females was quantified as the tally of her surviving F_3_ offspring (from a sample of 500 F_3_s). Male fitness was not estimated by Schluter et al. (2021), but the authors speculate that selection acted similarly on males and females because the evolutionary response observed was highly similar to what was expected from estimates of female fitness. Bay et al. (2017) estimated the number of mating events for Paxton Lake F_2_ benthic × limnetic hybrid females, and pure (i.e., pure) benthic and limentic males. Analysis of the data suggested that length–mating relationships were the same across groups (i.e., no significant interaction) so all data were grouped for analysis. Data were analyzed using simple linear models from which intercepts and slopes were extracted. Relationships were significant (in a Poisson generalized linear model) and positive in both datasets, but we chose to base fecundity estimates on the estimates from Schluter et al. (2021) because differences in composite fitness among groups were smaller (i.e., it is more conservative).

## Supplementary Tables & Figures

**Table S1.** Summary of fish numbers and survival (re-capture numbers) for the 2020 pond experiment.

| pond | origin | ecotype | type | *n* released | *n* recaptured | survival proportion |
| --- | --- | --- | --- | --- | --- | --- |
| 4 | Little Quarry | b | Pure | 102 | 46 | 0.45 |
| 4 | Paxton | b | Pure | 107 | 70 | 0.65 |
| 4 | hybrid | b | F<sub>1</sub> | 206 | 123 | 0.60 |
| 4 | hybrid | b | F<sub>2</sub> | 210 | 130 | 0.62 |
| 4 | Little Quarry | l | Pure | 106 | 31 | 0.29 |
| 4 | Paxton | l | Pure | 107 | 39 | 0.36 |
| 4 | hybrid | l | F<sub>1</sub> | 207 | 101 | 0.49 |
| 4 | hybrid | l | F<sub>2</sub> | 208 | 62 | 0.30 |
| 9 | Little Quarry | b | Pure | 104 | 84 | 0.81 |
| 9 | Priest | b | Pure | 106 | 93 | 0.88 |
| 9 | hybrid | b | F<sub>1</sub> | 203 | 185 | 0.91 |
| 9 | hybrid | b | F<sub>2</sub> | 205 | 169 | 0.82 |
| 9 | Little Quarry | l | Pure | 105 | 36 | 0.34 |
| 9 | Priest | l | Pure | 105 | 57 | 0.54 |
| 9 | hybrid | l | F<sub>1</sub> | 204 | 86 | 0.42 |
| 9 | hybrid | l | F<sub>2</sub> | 205 | 100 | 0.49 |
| 19 | Priest | b | Pure | 103 | 95 | 0.92 |
| 19 | Paxton | b | Pure | 103 | 87 | 0.84 |
| 19 | hybrid | b | F<sub>1</sub> | 204 | 174 | 0.85 |
| 19 | hybrid | b | F<sub>2</sub> | 205 | 161 | 0.79 |
| 19 | Priest | l | Pure | 105 | 32 | 0.30 |
| 19 | Paxton | l | Pure | 105 | 42 | 0.40 |
| 19 | hybrid | l | F<sub>1</sub> | 206 | 88 | 0.43 |
| 19 | hybrid | l | F<sub>2</sub> | 205 | 72 | 0.35 |

**Table S2.** Fitness components and composite fitness estimates for cross types within ecotype. We used estimates of the relationship between mass and standard length (Fig. S12), standard length and fecundity S13) and standard length and overwinter survival (Carlson et al., 2010) to generate estimates of composite fitness for each cross type for crosses of both ecotypes. Because the patterns were less conservative than a simple multiplicative estimate of composite fitness (‘comp. fit. mult.’ column), we opted to use the more conservative method.

| ecotype | cross type | summer surv. | mass (g) | std. lgth. (cm)* | comp. winter surv.* | fecundity* | comp. fit.† | comp. fit. mult. |
| --- | --- | --- | --- | --- | --- | --- | --- | --- |
| benthic | pure | 0.799 | 2.06 | 5.74 | 0.98 | 3.53 | 1 | 1 |
|  | F<sub>1</sub> | 0.818 | 2.14 | 5.84 | 1.00 | 3.69 | 1.08 | 1.07 |
|  | F<sub>2</sub> | 0.797 | 2.01 | 5.67 | 0.97 | 3.43 | 0.98 | 0.98 |
| limnetic | pure | 0.387 | 1.01 | 4.42 | 0.97 | 1.49 | 1 | 1 |
|  | F<sub>1</sub> | 0.454 | 1.09 | 4.56 | 1.00 | 1.69 | 1.34 | 1.27 |
|  | F<sub>2</sub> | 0.386 | 0.98 | 4.37 | 0.96 | 1.41 | 0.94 | 0.97 |

\* estimate; † $w = \text{sum. surv.} \times \text{wint. surv.} \times \text{fecundity}$

**Fig. S1.**
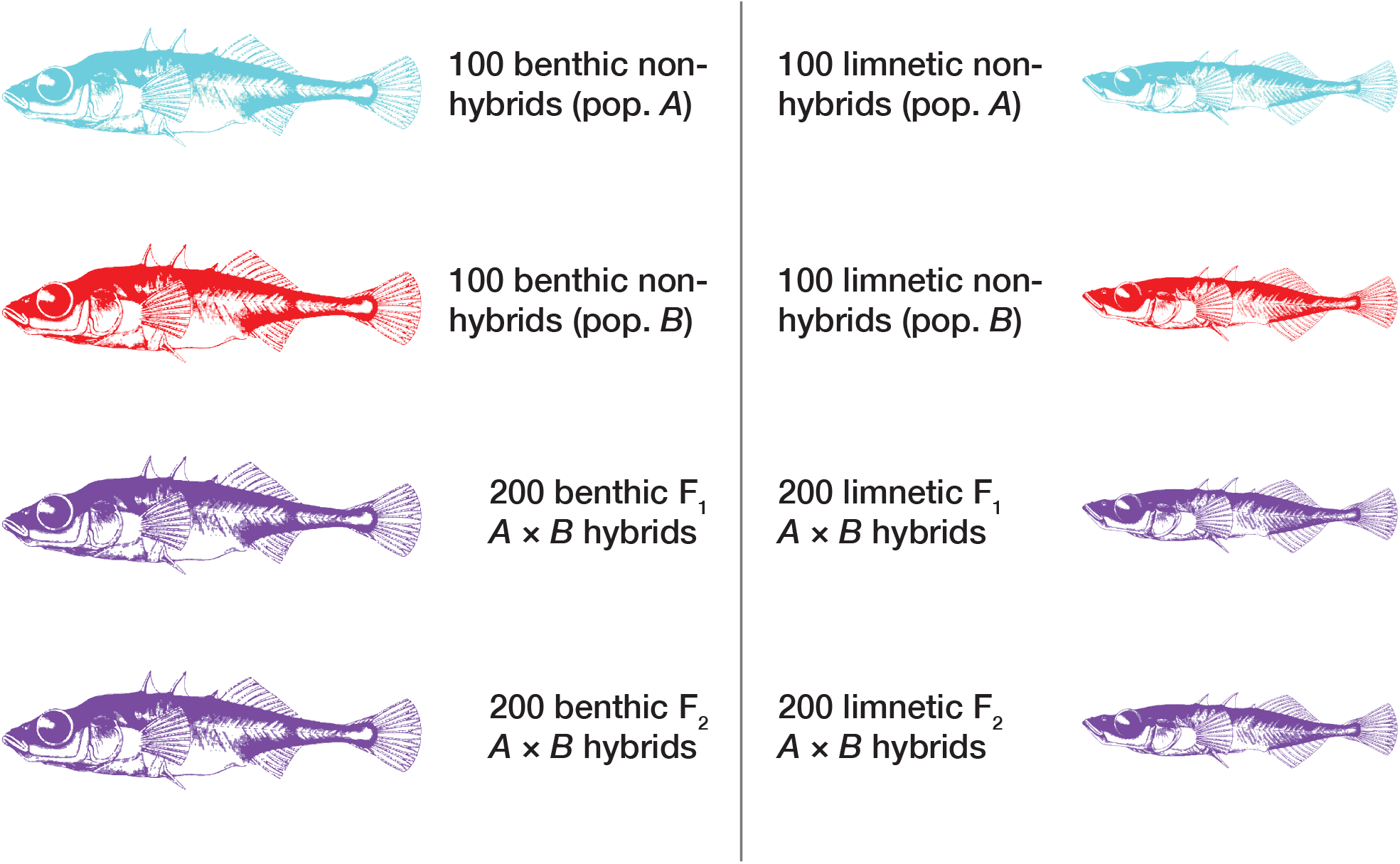
Overview of the experimental design. Each pond contained individuals from four cross types from each of the two ecotypes. In this figure (and not in the main text), red and cyan are the two different colours that represent two different pure populations (‘non-hybrids’). Purple represents (F_1_ and F_2_) hybrids.

**Fig. S2.**
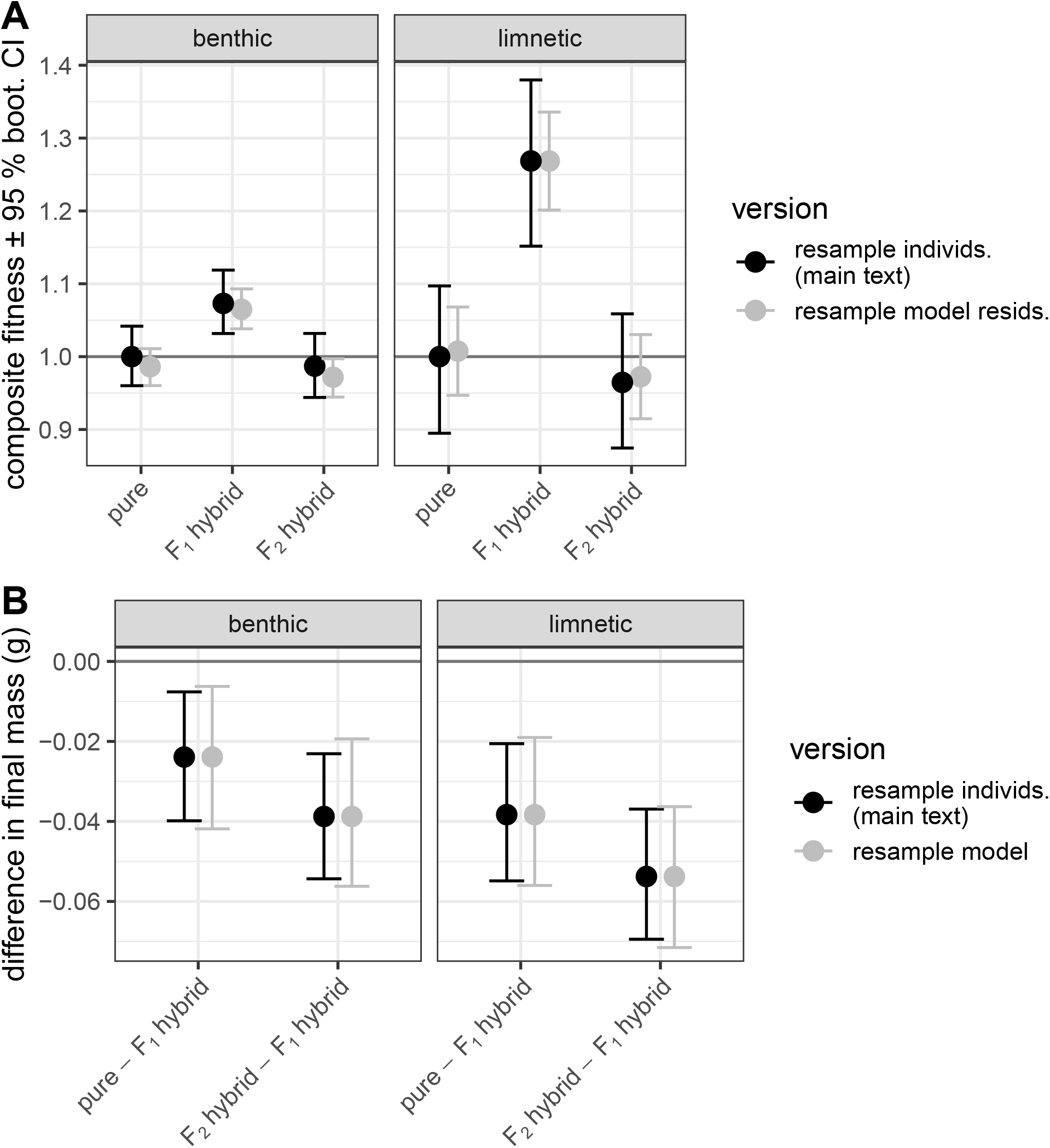
Conclusions of alternative bootstrap procedures. Panel (A) shows a case where model residuals are resampled (grey) in comparison to the resampling of individuals. Means are from the resampling but absolute fitness values are divided by the observed fitness of pure crosses. Panel (B) shows estimates of group differences in final mass (F_2_ – F_1_ and pure – F_1_) from a parametric bootstrap resampling procedure implemented via the `lmeresample` R package (Loy et al., 2021). This procedure has slightly wider confidence intervals than our approach because it simulates the error term for the ‘lake pair’ random effect, which with only three levels is estimated with poor accuracy.

**Fig. S3.**
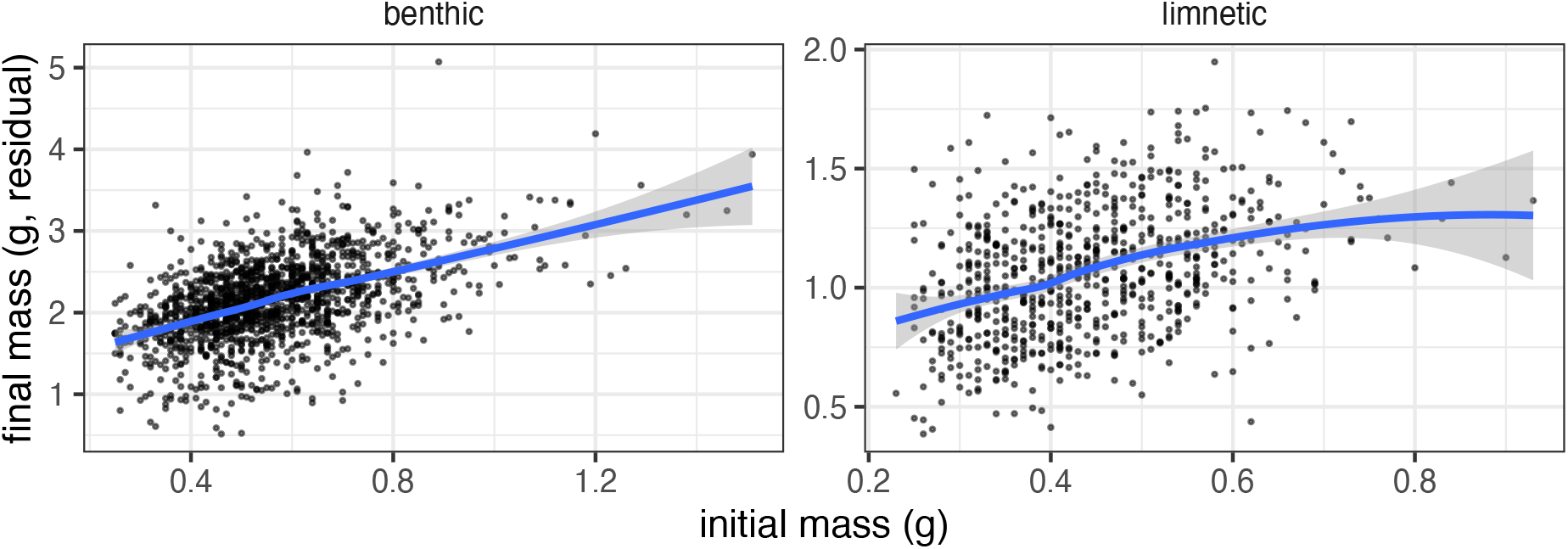
Relationship between initial and final mass in the pond experiment. The left plot shows benthic crosses and the right shows limnetic crosses. Points are partial residuals from a mixed model with lake pair as a random effect (pair-as-random models), extracted using visreg (Breheny and Burchett, 2017). We fit a smooth curve to the data here to illustrate patterns, but we analyze the mass data on a ln(mass + 1) scale.

**Fig. S4.**
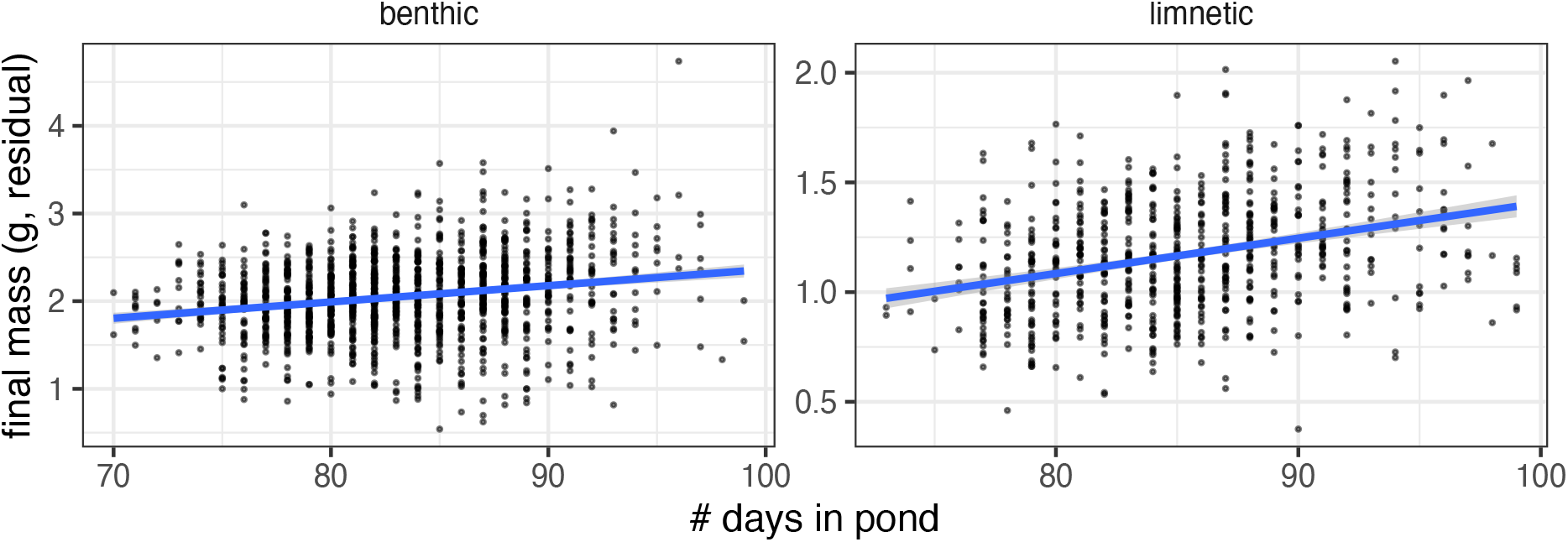
Relationship between the number of days in pond and final mass in the 2020 pond experiment. Both relationships are significant and positive. Note difference in axis scales.

**Fig. S5.**
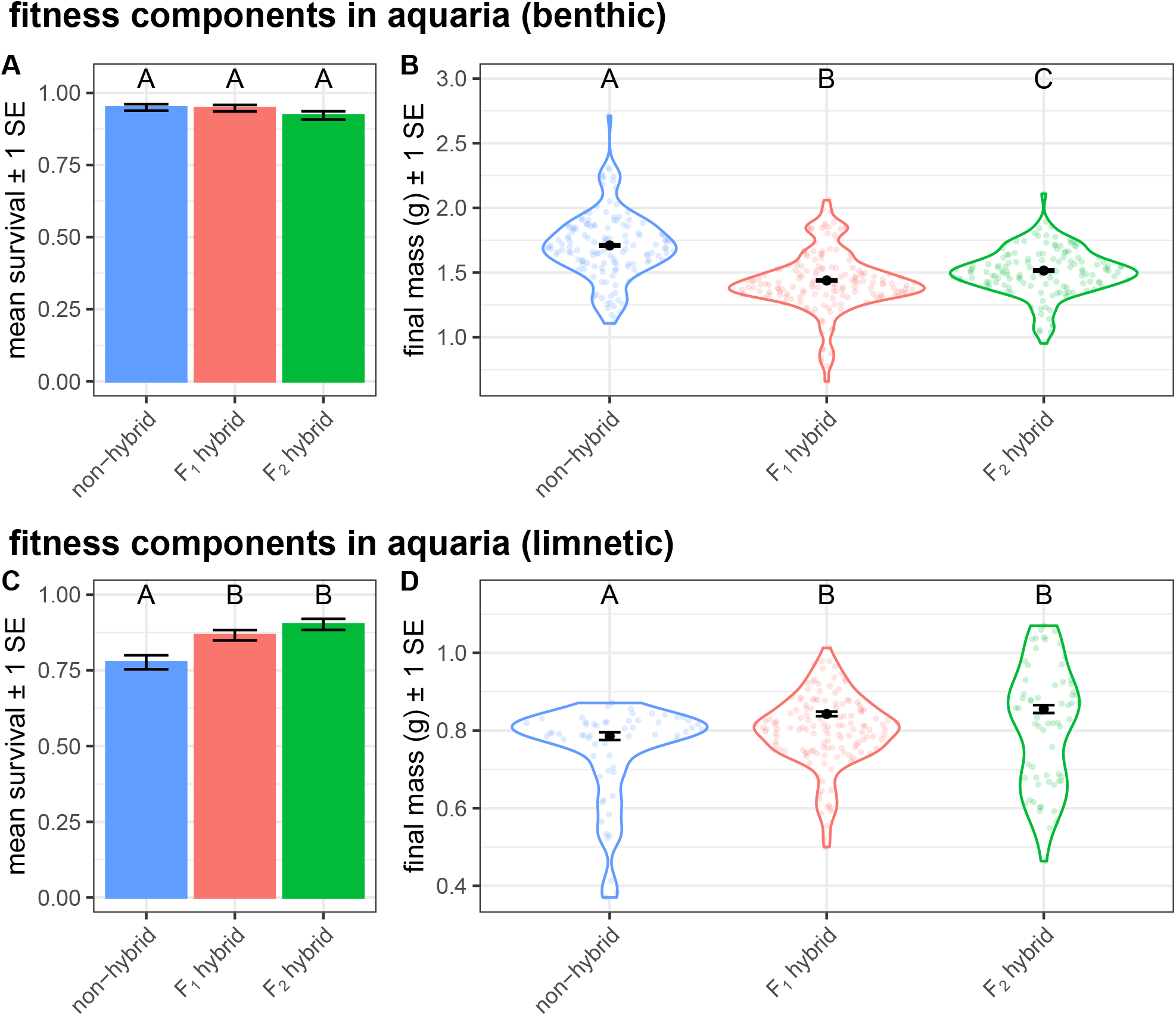
Components of fitness in aquaria. These data were used to generate the estimates of composite fitness in Fig. 3.

**Fig. S6.**
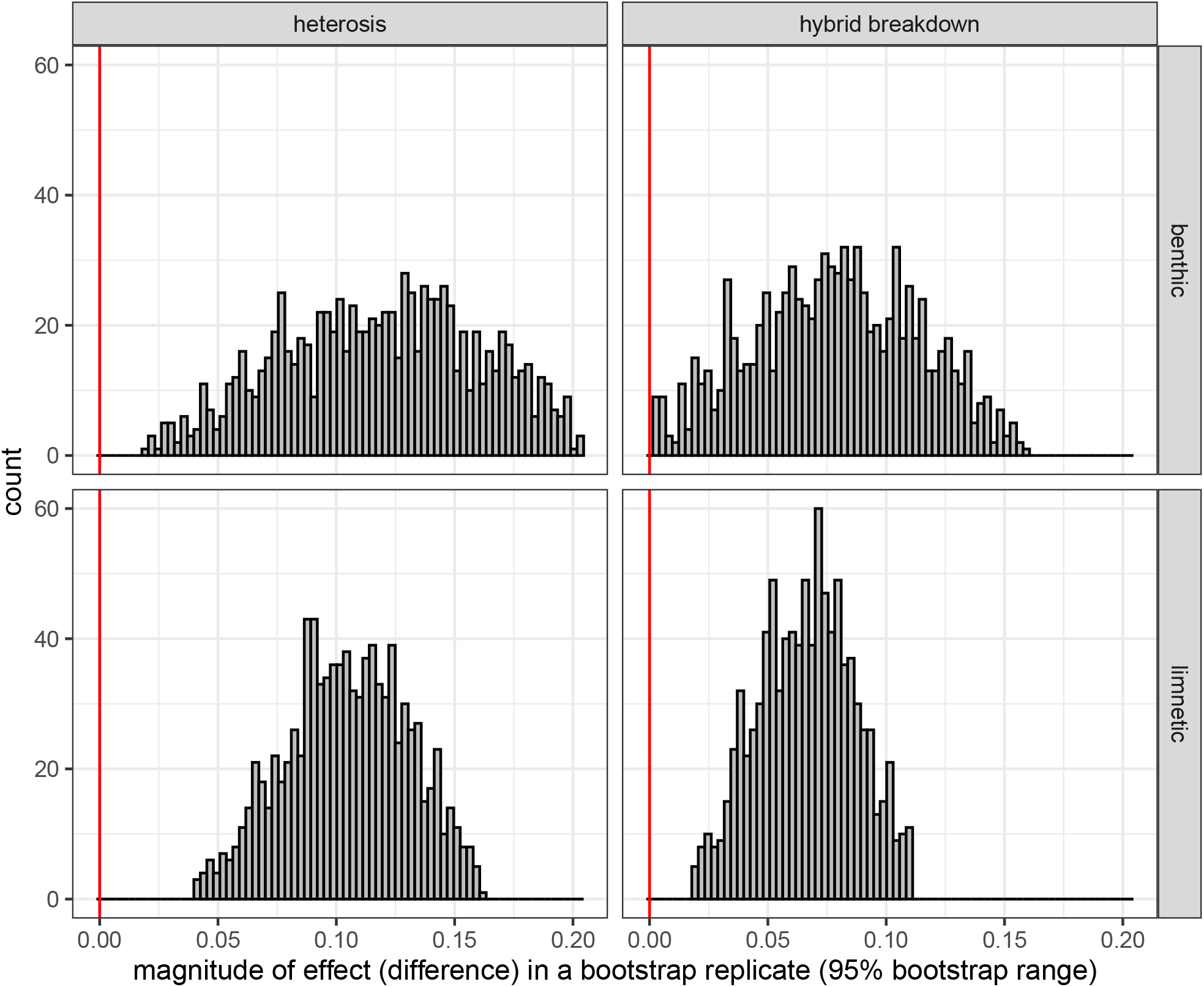
Distribution of bootstrap estimates of heterosis (left) and hybrid breakdown (right) via composite fitness for both benthic crosses (top) and limnetic crosses (bottom) (95% range). All 95% ranges do not include 0.

**Fig. S7.**
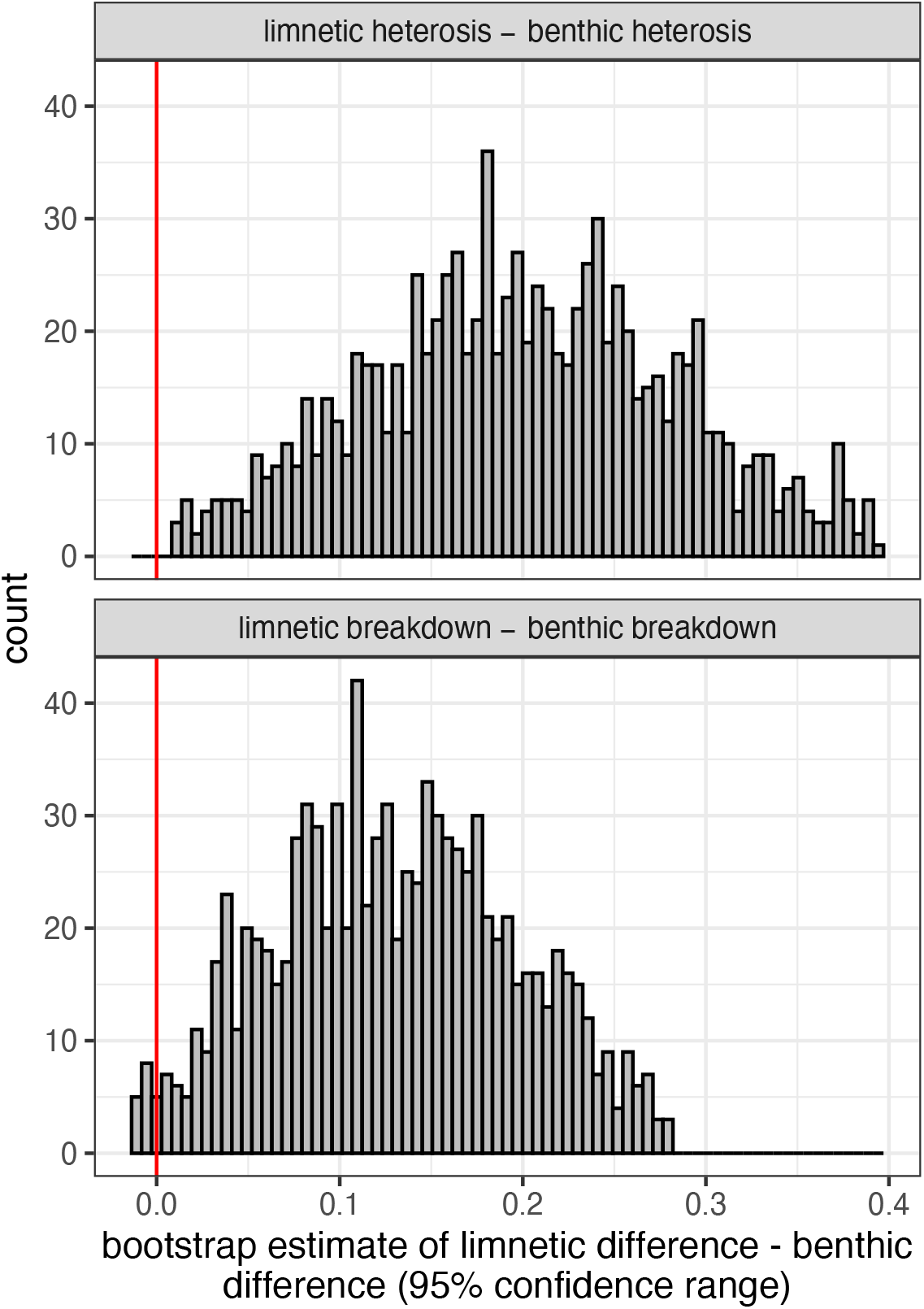
Comparison of the magnitude of heterosis and breakdown between benthic crosses and limnetic crosses. Heterosis and breakdown are both estimated as differences, and this analysis is therefore testing for a difference in these differences. Values show the 95% bootstrap confidence range, and ranges that do not include zero (red vertical line) are considered significant.

**Fig. S8.**
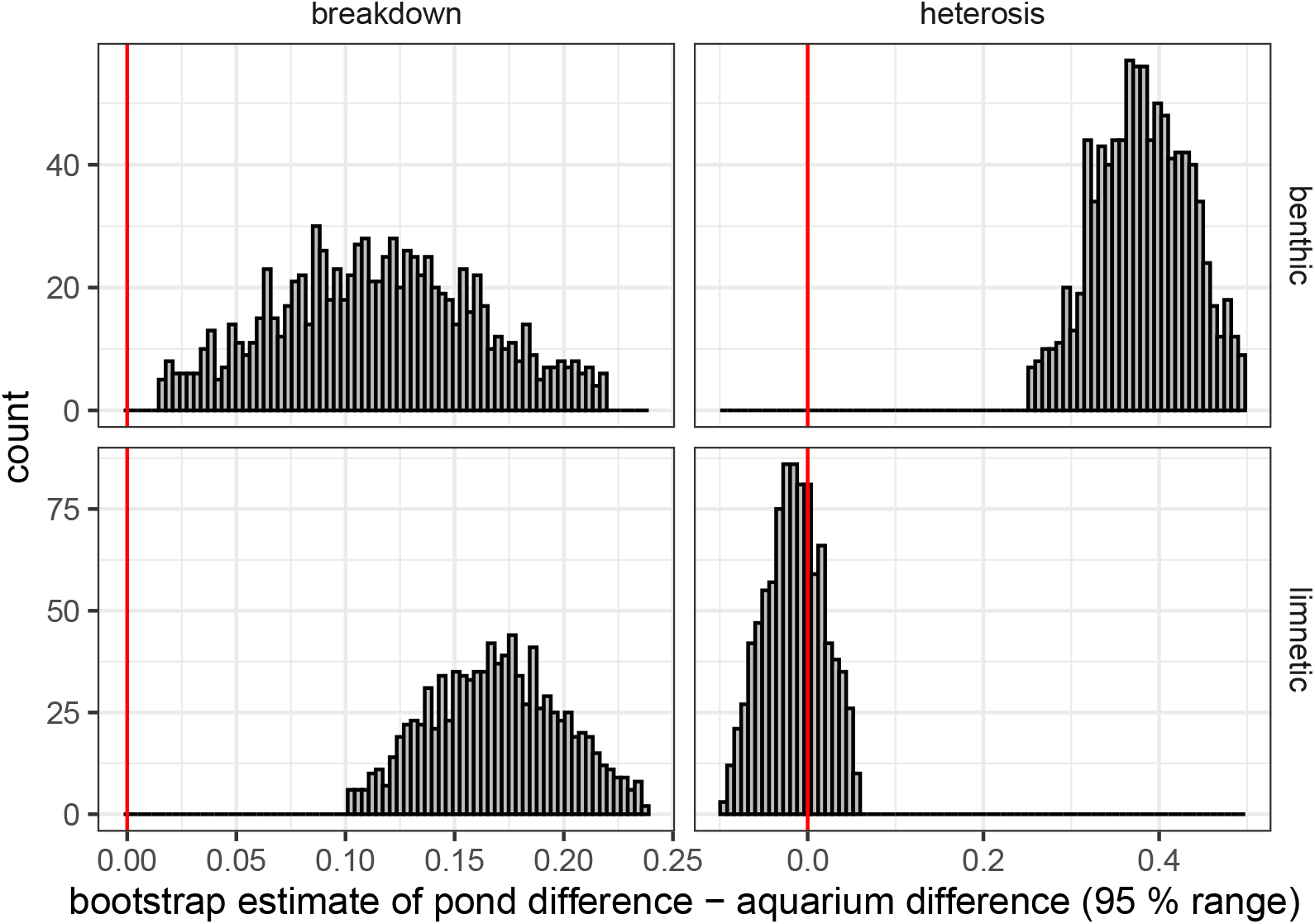
Comparison of the magnitude of heterosis and breakdown between pond and aquarium data (within ecotype). Heterosis and breakdown are both estimated as differences, and this analysis is therefore testing for a difference in these differences between the same metric in pond data vs. aquaria. Ranges that do not include 0 (red vertical line) are considered statistically significant.

**Fig. S9.**
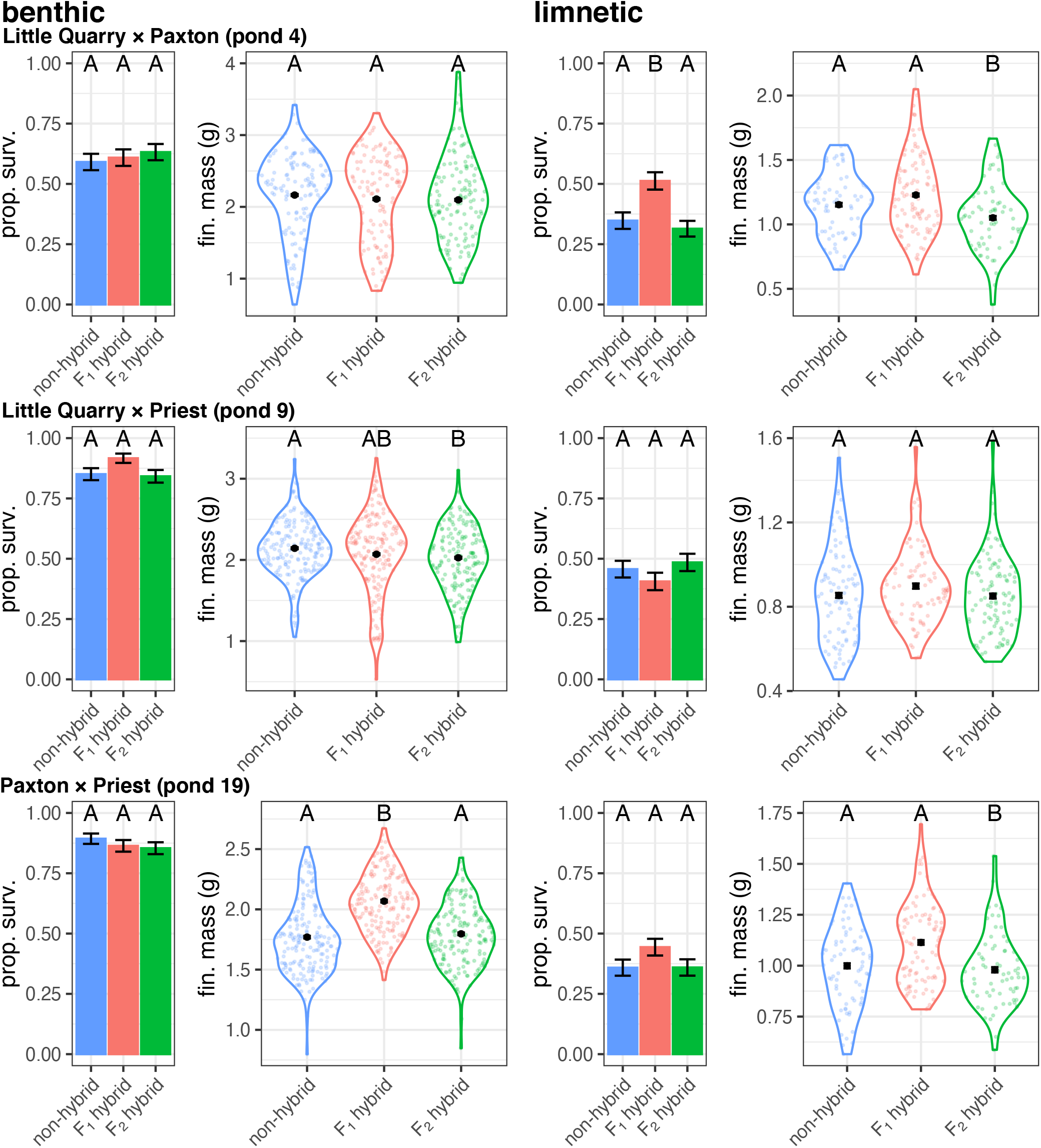
Components of fitness in each experimental pond (rows; see labels) for benthic crosses (left) and limnetic crosses (right). Error bars are ± 1SE.

**Fig. S10.**
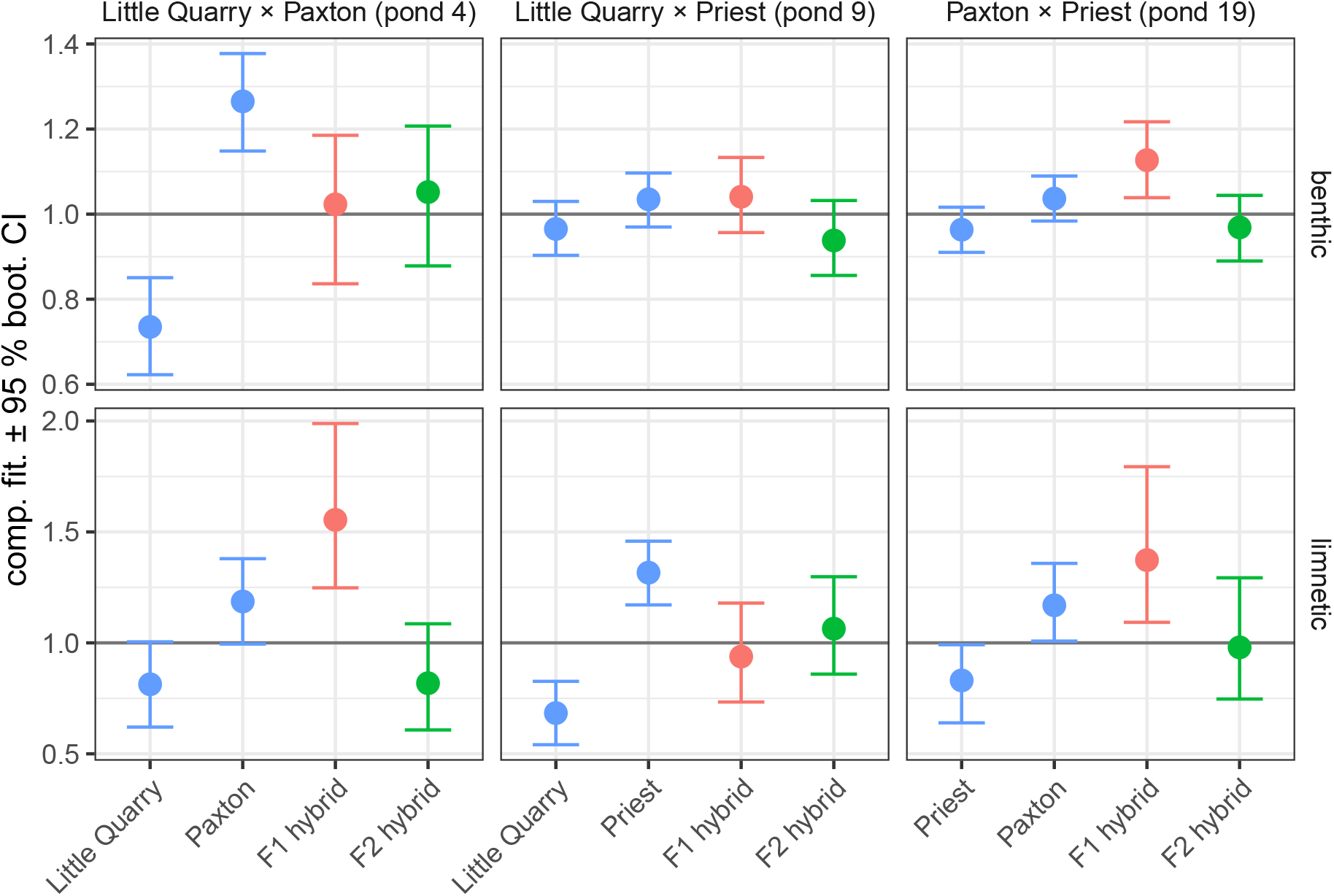
Fitness components with the two parent populations plotted separately. Points are estimated marginal means (emmeans package; Lenth et al. 2020), and 95% confidence intervals from 1,000 bootstrap replicates. In all cases where heterosis is observed, the F_1_ is either superior to both parents or significantly superior to the less fit parent with an overlapping interval (though higher mean) than the fitter parent.

**Fig. S11.**
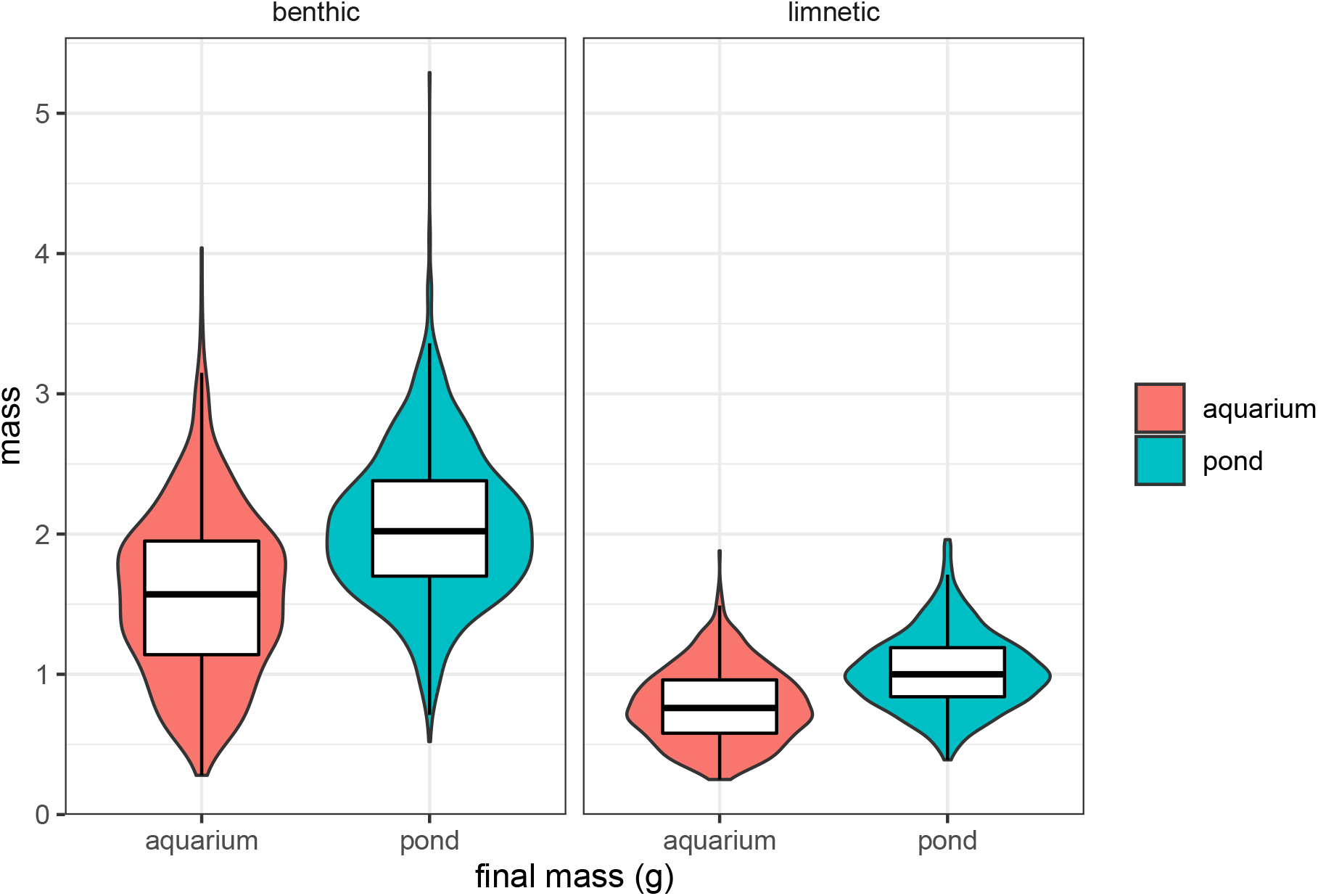
Final mass (g) of benthic (left) and limnetic (right) fish from aquarium (red) and pond (blue) experiments. Values are raw data, not from models. Fish from ponds were significantly larger than fish from aquaria for both ecotypes (both *P* < 0.0001).

**Fig. S12.**
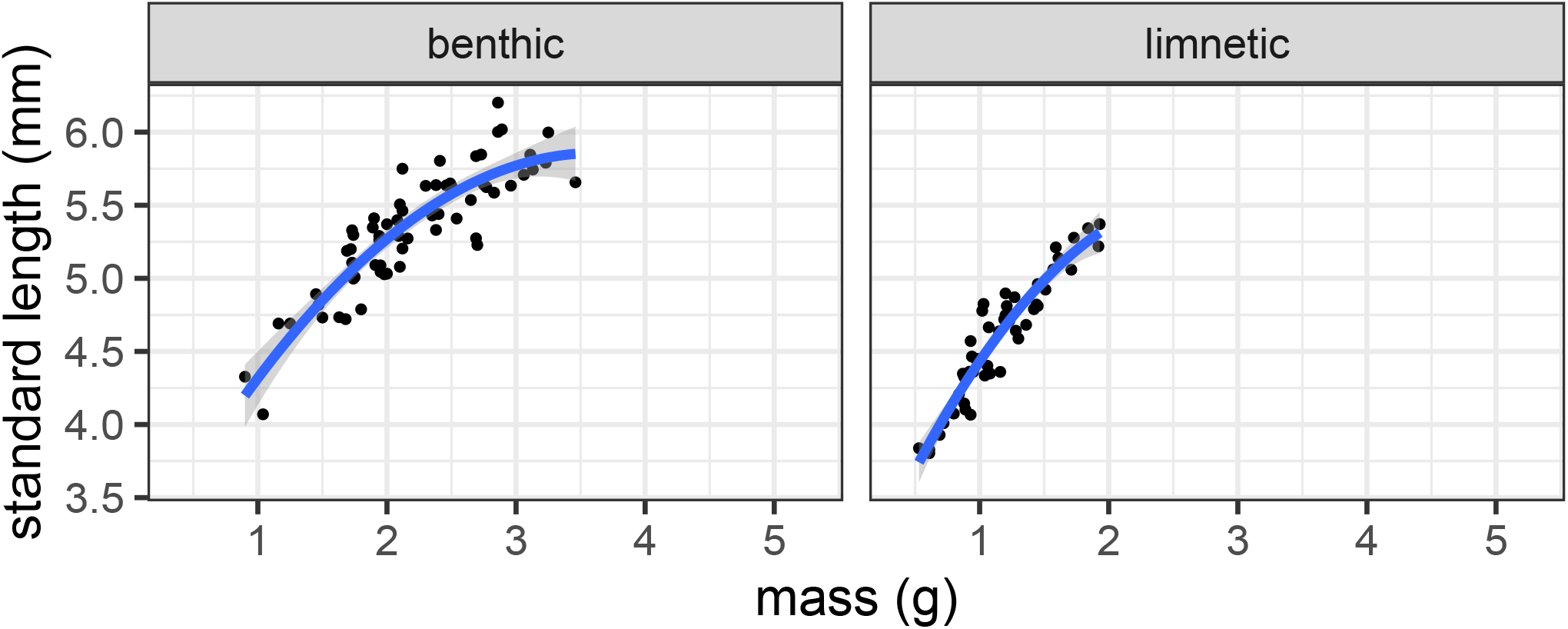
Relationship between mass and standard length for fish collected from ponds. Relationships are shown separately for benthic crosses and limnetic crosses. Both are significant and positive. These data were used to estimate standard length across the dataset.

**Fig. S13.**
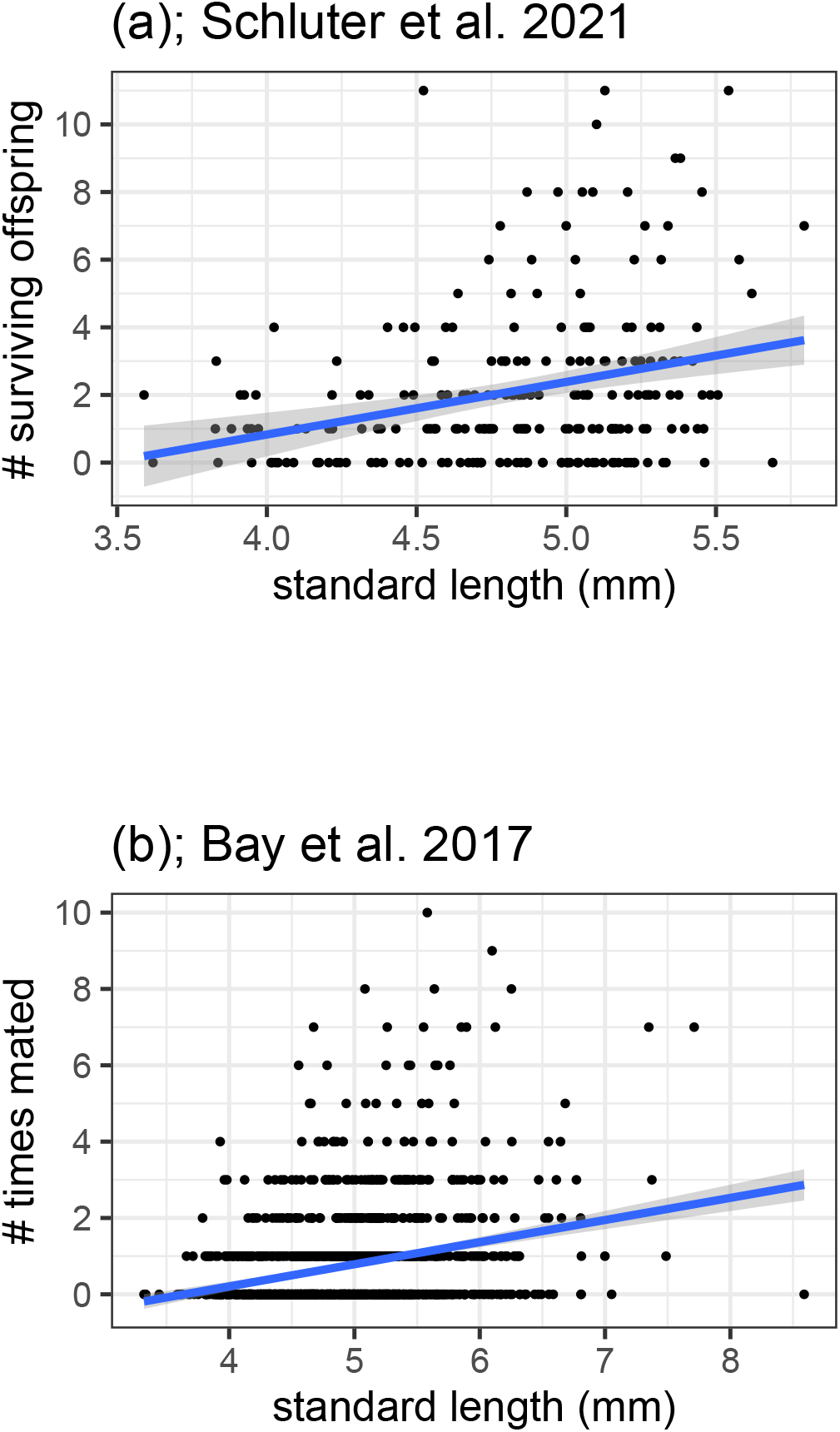
Relationship between standard length and fecundity in previously published stickleback pond experiments. Schluter et al. (2021) considered F_2_ marine-freshwater hybrid females and measured the number of F_3_ hybrids to which they could be confidently assigned parentage. Bay et al. (2017) considered F_2_ benthic-limnetic (Paxton) hybrid females and pure males of both ecotypes and recorded the number of successful mating events (inferred from genotyped eggs and offspring). A generalized linear model did not reject the hypothesis that the slope differed by group (F_2_ female, pure benthic male, or pure limnetic male), so we plot them together. Both relationships are significant and positive as evaluated with a generalized linear model.

